# Molecular glues that inhibit deubiquitylase activity and inflammatory signalling

**DOI:** 10.1101/2024.09.07.611787

**Authors:** Francesca Chandler, Poli Adi Narayana Reddy, Smita Bhutda, Rebecca L. Ross, Arindam Datta, Miriam Walden, Kieran Walker, Stefano Di Donato, Joel A. Cassel, Michael A. Prakesch, Ahmed Aman, Alessandro Datti, Lisa J. Campbell, Martina Foglizzo, Lillie Bell, Daniel N. Stein, James R. Ault, Rima S. Al-awar, Antonio N. Calabrese, Frank Sicheri, Francesco Del Galdo, Joseph M. Salvino, Roger A. Greenberg, Elton Zeqiraj

## Abstract

Deubiquitylases (DUBs) are crucial in cell signalling and are often regulated by interactions within protein complexes. The BRCC36 isopeptidase complex (BRISC) regulates inflammatory signalling by cleaving K63-linked polyubiquitin chains on Type I interferon receptors (IFNAR1). As a Zn^2+^-dependent JAMM/MPN DUB, BRCC36 is challenging to target with selective inhibitors. We discovered first-in-class inhibitors, termed <u>B</u>RISC molecular g<u>lues</u> (BLUEs), which stabilise a 16-subunit BRISC dimer in an autoinhibited conformation, blocking active sites and interactions with the targeting subunit SHMT2. This unique mode of action results in selective inhibition of BRISC over related complexes with the same catalytic subunit, splice variants and other JAMM/MPN DUBs. BLUE treatment reduced interferon-stimulated gene expression in cells containing wild type BRISC, and this effect was absent when using structure-guided, inhibitor-resistant BRISC mutants. Additionally, BLUEs increase IFNAR1 ubiquitylation and decrease IFNAR1 surface levels, offering a potential new strategy to mitigate Type I interferon-mediated diseases. Our approach also provides a template for designing selective inhibitors of large protein complexes by promoting, rather than blocking, protein-protein interactions.

## Introduction

There are over 100 human DUBs, which control cellular signalling by dictating protein activity, localisation, or stability^1–5^. DUB dysfunction is implicated in a range of pathologies, including autoimmune disorders, cancers, metabolic diseases, and neurodegeneration^6–8^. Consequently, DUBs remain attractive therapeutic targets and are the focus of many drug discovery efforts^9,10^.

BRCC36 is a JAMM (JAB1, MOV34, and MPR1, Pad1 N-terminal (MPN)) metalloenzyme DUB and selectively cleaves lysine 63-linked ubiquitin (K63-Ub) chains^11,12^. BRCC36 is present in two distinct macromolecular assemblies: a cytoplasmic BRCC36 isopeptidase complex (BRISC), and the nuclear Abraxas1 isopeptidase complex (ARISC). The BRISC complex regulates Type I interferon signalling by deubiquitylating and stabilising Type I interferon (IFNAR1) receptors, whilst the ARISC complex interacts with the tumour suppressor protein BRCA1 and localises to double-stranded DNA breaks to facilitate DNA damage repair^13–15^. BRCC36 (MPN^+^) requires a pseudo-DUB partner for enzymatic activity. This occurs through a stable heterodimeric complex with either Abraxas1 (MPN^-^) in the nucleus, or Abraxas2 (MPN^-^) in the cytoplasm^12,16–18^. BRISC and ARISC complexes contain two additional proteins, BRCC45 and MERIT40, and form dimer of hetero-tetramer assemblies with a 2:2:2:2 stoichiometry^16–19^. These eight subunit enzyme complexes require additional interacting partners for cellular function. BRISC forms a complex with a metabolic enzyme, serine hydroxymethyltransferase 2 (SHMT2), for targeting to IFNAR1 receptors, and loss of this interaction leads to a reduction in interferon signalling^20^ **(Extended Data Fig. 10**). ARISC forms the BRCA1-A complex with BRCA1, BARD1, and RAP80 in the nucleus to facilitate recruitment to DNA double strand breaks^13–15^. Recent cryo-electron microscopy (cryo-EM) structures of the BRISC-SHMT2 complex revealed a U-shaped assembly of the BRISC complex, whereby the BRCC36-Abraxas2 heterodimer bridges two BRCC45-MERIT40 “arms”^20,21^. A similar overall architecture was also observed for ARISC structures^19,21,22^.

BRISC-mediated deubiquitylation of IFNAR1 receptors promotes Janus kinase (JAK)/signal transducers and activators of transcription (STAT) signalling and expression of interferon (IFN) stimulated genes (ISGs)^23^. Elevated ISG expression is associated with autoimmune diseases, including Systemic Lupus Erythematosus (SLE)^24^, Rheumatoid Arthritis (RA)^25^, and Systemic Sclerosis (SSc)^26^. BRISC-deficient mice are protected from elevated interferon signalling and certain forms of inflammation^23^. Therefore, targeting BRISC with small molecule inhibitors represents a therapeutic strategy to reduce persistent inflammation and subsequent autoimmune disease driven pathology.

Significant progress has been made in selectively targeting the ubiquitin specific protease (USP) family DUBs^27–31^. These hold promise as potential therapeutics and as tool compounds to understand DUB biology. However, most inhibitors of the JAMM/MPN family of DUBs are broad-spectrum zinc chelators and there are currently no selective inhibitors for BRCC36 complexes^11,32^. Capzimin, a quinoline-8-thiol (8TQ) derivative targets the active site zinc of proteasomal subunit Rpn11, but also inhibits BRCC36 and AMSH^33^. Inhibitors of the JAMM domain containing de-neddylase, CSN5, also engage the catalytic zinc, but show specificity for CSN5 over AMSH and PSMD14^34^. Thus, whilst major progress has been made in DUB inhibitor development^35^, small-molecule inhibitors of the JAMM/MPN DUBs exclusively target the conserved zinc binding pocket, which makes the development of selective inhibitors challenging.

Molecular glues (MGs) are defined as small molecule stabilisers of protein-protein interactions^36,37^. Such compounds act as immunosuppressants (e.g. cyclosporin A^38,39^, rapamycin) and natural degraders (e.g. auxin in plants^37^). Immune-modulatory imide drugs (IMiDs), such as thalidomide, are MGs which induce protein degradation by stabilising an interaction between the E3 ligase cereblon and neo-substrates^40^. As such, MGs are an attractive class of compounds for regulating protein stability and degradation, however, MGs for DUBs have not been reported.

We describe first-in-class, selective BRISC inhibitors and define a unique mechanism of action for a DUB inhibitor. Cryo-EM structures of BRISC in complex with small molecule inhibitors reveal the molecular basis for selectivity and compound mechanism of action. The BRISC inhibitors identified here act as molecular glues and do not engage the active site zinc. Instead, <u>B</u>RISC molecular g<u>lues</u> (BLUEs) inhibit DUB activity by stabilising a BRISC conformer that occludes the BRCC36 active site from accepting ubiquitin chains for cleavage. We show target engagement in cells through structure-guided mutagenesis and cell-based studies, and we further validate inhibitor mechanism of action on human cells following IFN stimulation and from patients with aberrant IFNAR1 activation. Overall, this study showcases the therapeutic potential of MG compounds which promote specific protein-protein interactions to achieve selective inhibition of macromolecular complexes.

## Results

### Identification of first-in-class, selective BRISC inhibitors

We designed a biochemical screen to identify BRISC small-molecule inhibitors by measuring activity of a commercial K63-linked di-ubiquitin substrate with an internally quenched fluorophore (IQF) (**Fig. 1a**, *left*). Increased fluorescence was detected over time, enabling continuous readout of DUB activity (**Fig. 1a**, *right*). We screened an in-house compound library of 320 published and custom-made kinase inhibitors and identified compounds AT7519 (well H20) and YM201636 (well P12) as hits (**Fig. 1b**). Compound selectivity was assessed against the broad-spectrum ubiquitin specific peptidase 2 (USP2), and the serine protease trypsin, which cleave K63-Ub substrate under the same assay conditions. YM201636 (well P12) inhibited BRISC, trypsin, and USP2, suggesting this is a non-specific inhibitor, whilst what we presumed to be compound AT7519 (well H20) showed selective inhibition of BRISC DUB activity (**Extended Data Fig. 1a**). To further validate the hit compound in well H20, we purchased AT7519 from two commercial vendors, Synkinase and Selleckchem. Curiously, neither inhibited BRISC DUB activity in the IQF assay (**Extended Data Fig. 1b**). UV-Visible spectroscopy analyses showed a different spectrum for the compound in well H20 compared to the purchased AT7519 compounds (**Extended Data Fig. 1c**), suggesting the compound in well H20 was different to AT7519. Liquid chromatography-mass spectrometry (LC-MS) revealed the H20 compound was pure, with a mass of 555.55 Da instead of the expected mass of 382.25 Da^41^ (**Extended Data Fig. 1d**). This mass difference is consistent with the addition of a 2,6-dichlorobenzaldehyde group, which we reasoned could have been inadvertently added during chemical synthesis at either the piperidine or pyrazole ring. We synthesised two possible isomers: AP-5-144 and JMS-175-2 (**Fig. 1c**) and tested their inhibitory effects against BRISC. We found JMS-175-2 matched the profile of compound in well H20, inhibiting BRISC with an IC_50_ of 3.8 μM (**Fig. 1d**). Consistent with the JMS-175-2 structure, mass spectrometry fragmentation analyses showed that compound H20 contains the 2,6-dichlorobenzaldehyde modification at the pyrazole ring, and not the piperidine ring (**Extended Data Fig. 1e**). The AP-5-144 isomer did not inhibit BRISC DUB activity and using fragmentation analyses we confirmed AP-5-144 did not match the chemical structure of compound H20 (**Extended Data Fig. 1e**). These data confirm the chemical structure of the compound in well H20 and led to the serendipitous identification of the BRISC inhibitor JMS-175-2.

**Figure 1.**
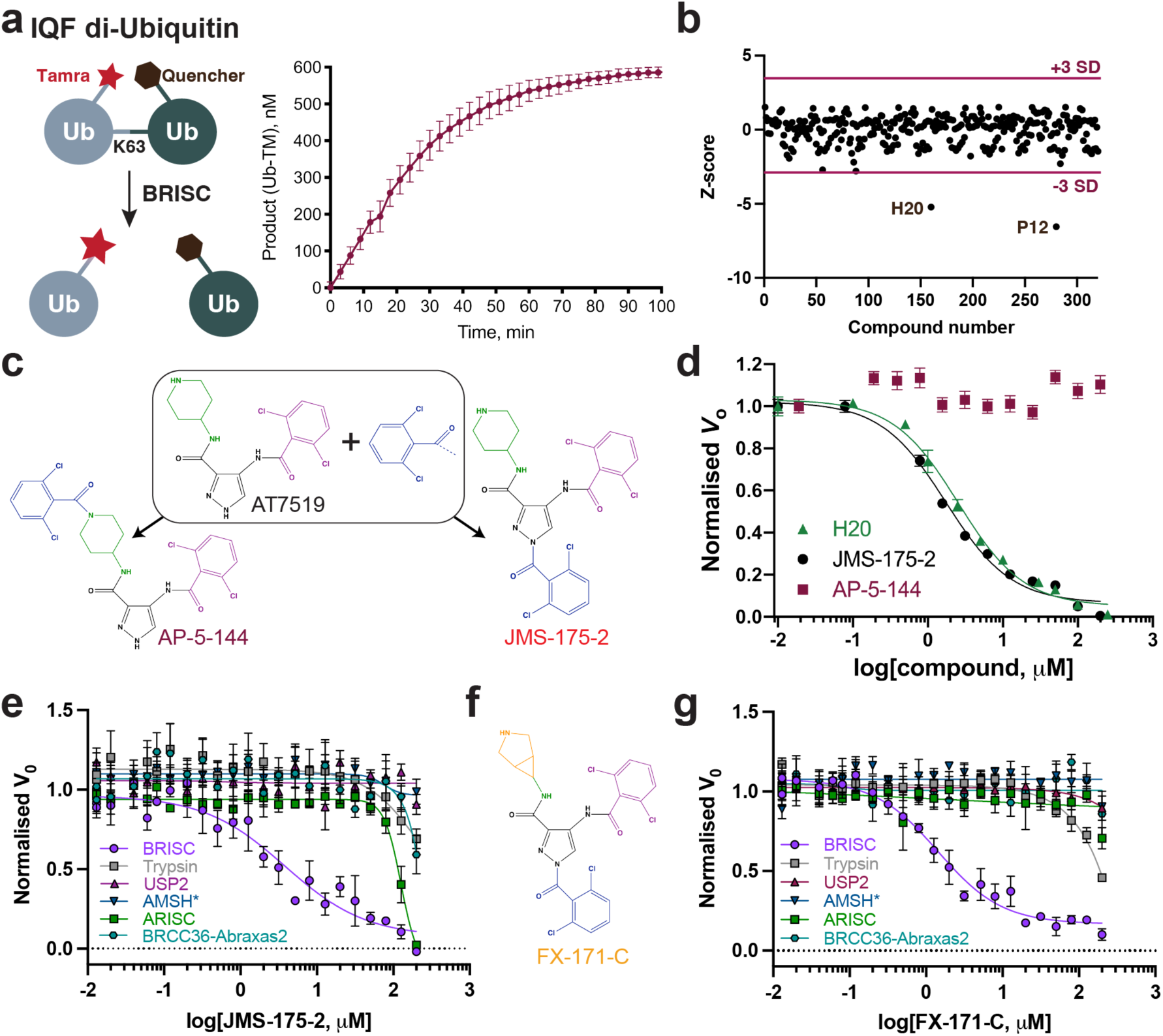
Fluorescence-based screen to identify first-in-class JAMM inhibitors. **a,** Schematic of a TAMRA-linked internally quenched fluorescent (IQF) di-ubiquitin substrate (*left*) and reaction progress curve of BRISC DUB activity (*right*). **b,** Z-score normalisation of 320 compounds from an in-house kinase-directed inhibitor library and identification of hit compounds in wells H20 and P12. SD = standard deviation. **c,** Chemical structures of AT7519 and of two isomers with an additional 2,6-dichlorobenzaldehyde moiety. **d,** Dose-response inhibition of BRISC activity by the H20 compound and the two potential isomers, AP-5-144 and JMS-175-2. **e, g,** Dose-response inhibition of trypsin, USP2 and JAMM/MPN DUB enzymes AMSH* (a STAM2-AMSH fusion^42^), BRISC, ARISC, and BRCC36-Abraxas2 by the indicated compounds. Data points in **d, e,** and **g,** are mean ± SEM of three independent experiments carried out in technical duplicate. **f,** Chemical structure of the FX-171-C compound.

We next determined the selectivity of inhibitors for the BRISC DUB beyond USP2 and trypsin. AMSH is a related JAMM/MPN DUB which, like BRCC36, selectively cleaves K63-linked polyubiquitin chains^43,44^. JMS-175-2 did not inhibit AMSH* (a STAM2-AMSH fusion)^42^ (**Fig. 1e**), showing it is selective for BRISC over other zinc-dependent DUBs. Remarkably, JMS-175-2 did not inhibit the nuclear ARISC complex, which shares three of the four BRISC subunits, including the catalytic subunit BRCC36 (**Fig. 1e**). A related analogue, FX-171-C (**Fig. 1f**), had a moderately improved IC_50_ of 1.4 μM compared to JMS-175-2 (IC_50_ = 3.8 μM), and retained selectivity for BRISC against other JAMM/MPN DUBs (**Fig. 1g**). These data confirm JMS-175-2 series are selective BRISC inhibitors and suggest the specificity is conferred, in part, by the Abraxas2 subunit which is substituted for Abraxas1 in the ARISC complex.

To fully explore the selectivity profile of JMS-175-2, we evaluated its inhibitory effects on 48 DUBs spanning five DUB families. JMS-175-2 did not fully inhibit any of the DUBs present in the panel, including AMSH-LP (**Extended Data Fig. 1f**).

Curiously, JMS-175-2 and FX-171-C did not inhibit the minimally active BRCC36-Abraxas2 complex, which indicates that the “arm” regions containing BRCC45 and MERIT40 also contribute to the inhibitor selectivity profile (**Figs. 1e, 1g**). We noticed a biphasic mode of inhibition (**Extended Data Fig. 2a**), and enzyme activity inhibition plots at different substrate concentrations suggested JMS-175-2 and FX-171-C act as noncompetitive inhibitors (**Extended Data Fig. 2b**). The strong selectivity of the JMS-175-2 and FX-171-C compounds and a noncompetitive mode of inhibition indicate these inhibitors do not target the Zn^2+^ active site, unlike previously described JAMM/MPN inhibitors^32–34^.

We used a commercially available fluorescently labelled K63-linked tetraubiquitin substrate to demonstrate both JMS-175-2 and FX-171-C inhibit BRISC-mediated cleavage of polyubiquitin chains (**Extended Data Fig. 2c**). The other possible JMS-175-2 stereoisomer, AP-5-144, did not inhibit BRISC cleavage of tetraubiquitin chains, consistent with the di-ubiquitin fluorescence assay (**Fig. 1d**). Importantly, JMS-175-2 and FX-171-C did not inhibit ARISC activity against tetraubiquitin (**Extended Data Fig. 2c**).

These experiments identify the first selective BRISC inhibitors and suggest a unique mechanism of action whereby the Abraxas2 pseudo-DUB subunit, and the BRCC45-MERIT40 “arms” contribute to selective inhibition.

### BRISC inhibitors stabilise an autoinhibited dimer conformation

To understand the molecular basis of BRISC inhibition by the new inhibitor series, and to determine the small molecule binding site, we characterised the complex by mass photometry and cryo-EM. Single molecule mass photometry measurements in the absence of any inhibitors revealed three populations of purified BRISC complexes. The major population corresponded to a single BRISC complex with four subunits at a 2:2:2:2 ratio (**Fig. 2a**, *top*), consistent with negative stain EM 2D class averages (**Fig. 2a**, *top inset*) and with previous studies^17,19,20^. We also observed a population at 163 kDa which may correspond to a dissociated 1:1:1:1 complex, or the BRCC36-Abraxas2 super dimer and minimally active complex^18^. Surprisingly, we also observed a third population, consisting of 2-5% of the particles, with an estimated molecular weight of 664 kDa. This corresponds to the mass of two BRISC “monomer” complexes with a predicted 4:4:4:4 stoichiometry.

**Figure 2.**
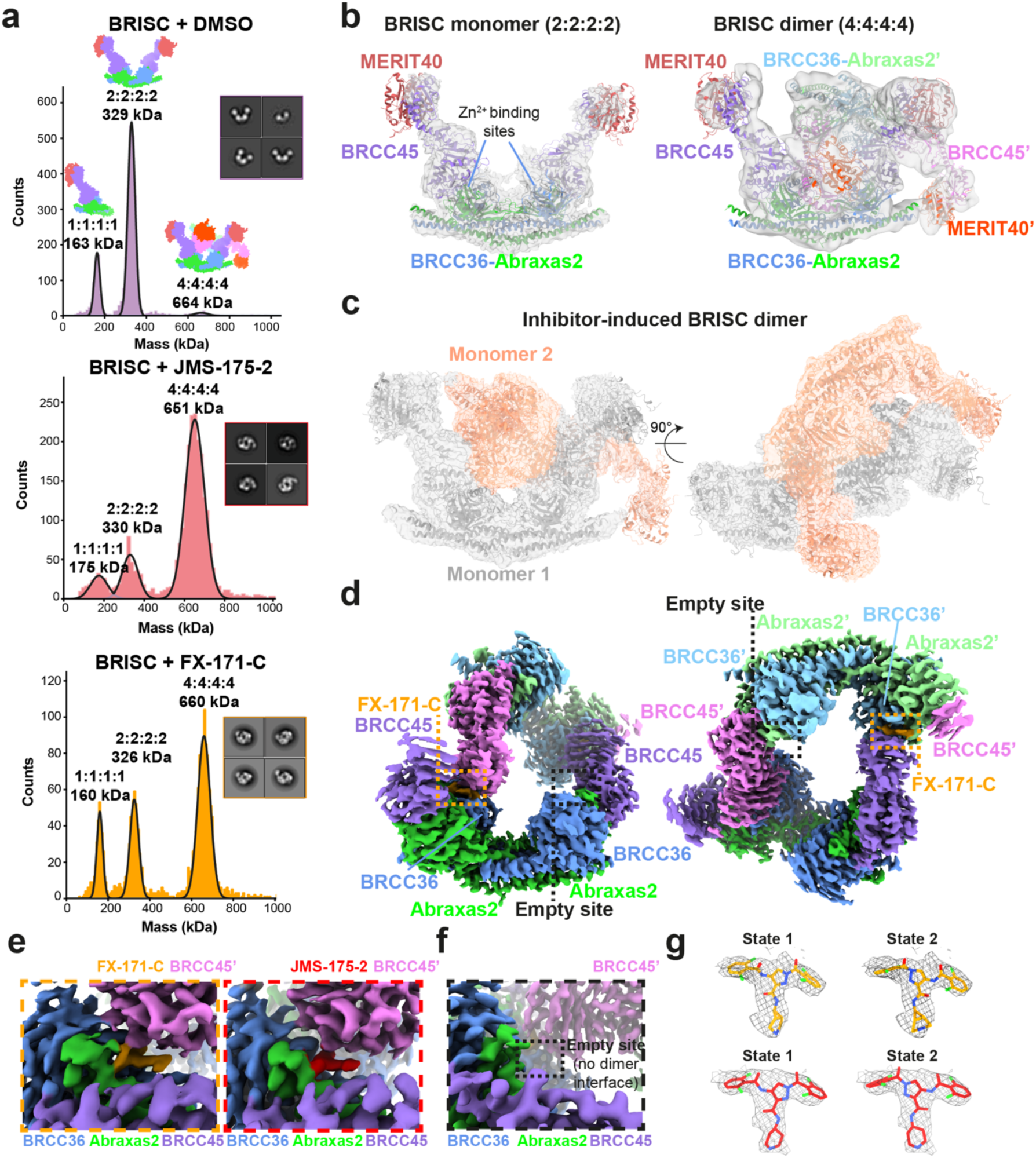
Inhibitors stabilise an inhibited BRISC dimer. **a,** Mass photometry histograms of purified BRISC in absence (DMSO, *top*) and presence of inhibitors (JMS-175-2, *middle*; FX-171-C, *bottom*), and corresponding negative stain EM 2D classes of BRISC mixed with DMSO or inhibitors (*insets*). **b,** *Left,* cryo-EM density map of a BRISC monomer with BRISC model (PDB: 6H3C) rigid-body fitted (dust cleaning size 7.4, map threshold 0.0907). *Right,* cryo-EM density map of a BRISC dimer with two BRISC models rigid-body fitted. Maps are outputs from non-uniform refinement in cryoSPARC. **c,** Cryo-EM density map of BRISC-FX-171-C co-structure at 3.0 Å. BRISC monomers are shown as grey and salmon cartoon models and fitted to the cryo-EM map shown as a transparent surface at 0.00224 threshold. The C-termini of BRCC45 (residues 275-383) and MERIT40 are rigid-body fitted into the density. **d,** BRISC-FX-171-C cryo-EM density map at 0.0165 threshold. BRISC subunits are coloured by chain. The density corresponding to FX-171-C is coloured orange and highlighted in orange boxes. The map shown in **c,** and **d,** is a locally-filtered map generated using RELION local resolution estimation. **e,** Close-up views of the indicated inhibitor density comparing FX-171-C (*left*) and JMS-175-2 (*right*) binding sites. **f,** Cryo-EM density at the equivalent sites of BRCC36, Abraxas2, and BRCC45’ in the BRISC-FX-171-C co-structure where there is no dimer interface, and no additional density corresponding to FX-171-C. The maps in **d-f** had dust cleaning (size 7.1) applied in ChimeraX. **g,** Structures of FX-171-C and JMS-175-2 modelled in State 1 and State 2. Cryo-EM density of the inhbitor after focused refinement represented as a mesh and displayed using the surface zone tool (FX-171-C radius 2.6, JMS-175-2 radius 2.2) in ChimeraX.

Consistent with these measurements, we observed a higher molecular weight BRISC species in cryo-EM data. We observed both BRISC “monomer” and BRISC “dimer” complexes in 2D class averages and in maps from *ab initio* reconstruction (**Extended Data Figs. 2d, 2e**). The majority of particles correspond to a monomeric complex, consistent with the BRISC-SHMT2 structure^20,21^ (**Fig. 2b, Extended Data Figs. 2e, 2f, Extended Data Table 1**). In addition, a low resolution cryo-EM reconstruction of particles corresponding to the BRISC dimer indeed contains density for two BRISC molecules (**Fig. 2b, Extended Data Fig. 2g, Extended Data Table 1**). The conformation of this dimeric BRISC species was different from the symmetric BRISC and ARISC dimers previously reported in glutaraldehyde cross-linked samples imaged by negative stain EM^19,21^ (**Extended Data Fig. 2h**). These observations suggest that BRISC has a propensity to dimerise, raising the possibility that these low-level dimers may be regulated or stabilised by ligand binding.

Interestingly, incubating purified BRISC with JMS-175-2 and FX-171-C resulted in a considerable mass shift to the 4:4:4:4 complex, which suggests the inhibitor promotes BRISC dimer formation (**Fig. 2a**, *middle and bottom*). Negative stain EM confirmed the oligomeric state on inhibitor addition, with 2D class averages that look like two U-shaped BRISC assemblies (**Fig. 2a**, *middle and bottom insets*). Using native mass spectrometry we confirmed the inhibitor-induced mass corresponds to a dimeric BRISC complex and BRISC dimers with 4:4:4:4 stoichiometry were detected after addition of JMS-175-2 and FX-171-C compounds (**Extended Data Figs. 3a, 3b**). Importantly, we also observed a dose-dependent increase in dimer formation by mass photometry for both JMS-175-2 and FX-171-C (**Extended Data Fig. 3c**). These data suggest an unexpected mode of action where inhibitor binding promotes a stable BRISC dimer complex of 16 subunits and molecular weight of 655 kDa.

### Cryo-EM structures reveal BRISC inhibitors act as molecular glues

To determine the precise mechanism by which a small molecule can induce formation of a multimeric DUB complex, we solved two co-structures of BRISC complexes, bound to FX-171-C and JMS-175-2. We observed a high proportion (>95%) of BRISC dimers after incubation with each inhibitor. After 3D refinement and postprocessing, we obtained cryo-EM maps at 3.0 Å (FX-171-C) and 3.3 Å (JMS-175-2) resolution (**Fig. 2c, Extended Data Figs. 4a-f, Extended Data Table 1**). The BRISC-inhibitor structures consist of a BRISC dimer, with density for all 16 subunits (stoichiometry 4:4:4:4), where the BRCC45-MERIT40 “arms” of one BRISC monomer hooks around the BRCC45-MERIT40 arm of a neighbouring BRISC molecule (BRISC’), bridging the BRCC36-Abraxas2 super dimer (**Fig. 2c**). Modelling of K63-linked di-ubiquitin substrate in this conformation suggests that the recruitment of a second BRISC octamer occludes the BRCC36 active sites by sterically blocking chain binding and catalysis (**Extended Data Fig. 3d**).

We observe the highest resolution (2.8-3.6 Å) in the core of the BRISC dimer structures, consisting of the BRCC36-Abraxas2 super dimer and the BRCC45’ subunit which forms the dimer interface (**Fig. 2d, Extended Data Figs. 4c, 4f**). The resolution is lower (7-12 Å) for the extreme C-termini of BRCC45 and MERIT40 (arm regions), therefore limiting accurate model building of these regions. This is due to the flexible nature of the arm regions and is consistent with our previous observations of the BRISC-SHMT2 cryo-EM structure^20^. Due to the lower resolution of the map beyond the second ubiquitin E2 variant (UEV) domain of BRCC45 and for MERIT40, we rigid-body fitted BRCC45 UEV-C (residues 275-383) and MERIT40 from previous BRISC-SHMT2 structures^20,21^.

The binding interface formed by BRCC36, Abraxas2, and BRCC45’ is also formed at the opposite site of the dimer structure (BRCC36’, Abraxas2’, BRCC45). At both interfaces, we observe additional density which is not attributed to either BRISC monomer and the density has the size and shape expected for each inhibitor (**Fig. 2e**). Importantly, the equivalent BRCC36-Abraxas2 surface that is not in contact with BRCC45’ from an opposing BRISC monomer does not contain additional cryo-EM density (**Fig. 2f**). The extra density is present in the same location for both the FX-171-C and JMS-175-2 maps (**Fig. 2e**), indicating a similar mode of binding for both compounds. Due to the slight tilting of the BRISC’ monomer resulting in an asymmetric dimer, there is one inhibitor bound per BRISC molecule, and two inhibitors per BRISC dimer (4:4:4:4:2 stoichiometry).

Focused refinement using a mask comprising the core of the BRISC dimer moderately improved the density for the FX-171-C compound (**Extended Data Figs. 4g, 4h**). Likewise, applying a mask on the highest resolution half of the JMS-175-2 map also improved the density for JMS-175-2 (**Extended Data Figs. 4i, 4j**). Due to the presence of two dichlorobenzene rings in each compound, we were unable to unambiguously determine the orientation of the dichlorobenzene moieties in the cryo-EM densities and have modelled the ligands in two orientations: State 1 and State 2 (**Fig. 2g**).

Next, we examined the conformational changes induced by FX-171-C using differential hydrogen deuterium exchange-mass spectrometry (HDX-MS) analysis. Measuring differences in deuterium uptake, detected at the peptide level, in the absence and presence of FX-171-C enabled us to analyse the structural rearrangement after inhibitor binding (**Extended Data Fig. 5a**). For example, regions of protection upon FX-171-C addition were identified in BRCC36 (residues 111-135) and BRCC45 (residues 122-134), which are consistent with the small molecule binding site and interaction interfaces identified in our cryo-EM structures (**Extended Data Figs. 5b, 5c**). We also observed deprotection of a BRCC36 peptide (142-149), indicative of a change in solvent accessibility near the enzyme active site. Protected peptides in BRCC45 (residues 206-221, 311-327) suggest further interactions between BRCC45 subunits from opposing BRISC monomers (**Extended Data Fig. 5b**). Moreover, deprotected peptides in the BRCC36-Abraxas2 coiled-coil and the C-termini of BRCC45 and MERIT40 subunits indicate additional and far-reaching conformational changes induced by inhibitor binding (**Extended Data Fig. 5b**).

Collectively, these structural analyses establish the inhibitors are <u>B</u>RISC molecular g<u>lues</u> (BLUEs) which stabilise two BRISC octamers to form a BRISC dimer with 16-subunits. BLUEs bind at a composite site of three interacting proteins: BRCC36 and Abraxas2 from one BRISC monomer and BRCC45’ from a second BRISC monomer. The inhibitor-induced dimer is an inactive conformation, whereby ubiquitin chain binding and processing is blocked.

### The BLUE binding pocket

The BLUE compound binding pocket is in close proximity to, but does not engage, the catalytic zinc and does not interact with BRCC36 active site residues (**Extended Data Fig. 5d**), consistent with enzyme activity data suggesting that BLUEs are non-competitive inhibitors (**Extended Data Fig. 2b**). BLUE compounds are the first examples of non-competitive JAMM/MPN DUB inhibitors, as all known JAMM/MPN DUB inhibitors described to date target the zinc binding site^32–34,45^ (**Extended Data Fig. 5e**). In addition to exploring a new binding site for JAMM/MPN DUBs, BLUE compound engagement of the middle BRCC45 ubiquitin E2 variant (UEV) domain highlights another unexpected compound binding surface in E2 folds. Unlike BAY 11-7082 and NSC697923 (inhibitors of Ubc13), BLUEs do not engage the UEV pseudo-catalytic site^46^, nor bind to an allosteric site exemplified by the Cdc34 E2 inhibitor, CC0651^47^ (**Extended Data Fig. 5f**).

The local resolution of our cryo-EM structures at the dimer interface is ∼2.8 Å (FX-171-C map, (**Fig. 2e**)) and sufficient to identify residues from each BRISC monomer which contribute to the inhibitor binding pocket. In BRCC36, BLUEs bind between the S-loop (*β*4-*α*3) and the *β*5, *β*6-strands. In Abraxas2, BLUE compounds interact with the *β*5-strand and *β*5-*β*6 loop (**Fig. 3a**, *left*). Two *α*-helices (*α*6 and *α*10) from the BRCC45’ subunit also line the inhibitor binding pocket (**Fig. 3a**, *right*). The two dichlorobenzene moieties of JMS-175-2 and FX-171-C sit in a hydrophobic groove formed by BRCC36 S-loop residues T128 and W130, and residues I158 and L169 (**Fig. 3b**). Abraxas2 I133 and BRCC45’ F140, C245, and I247 also contribute to the hydrophobic binding pocket. The JMS-175-2 piperidine ring and FX-171-C pyrrolidine ring extend into a hydrophilic region encompassing BRCC36 D160 and R167, and BRCC45’ D248.

**Figure 3.**
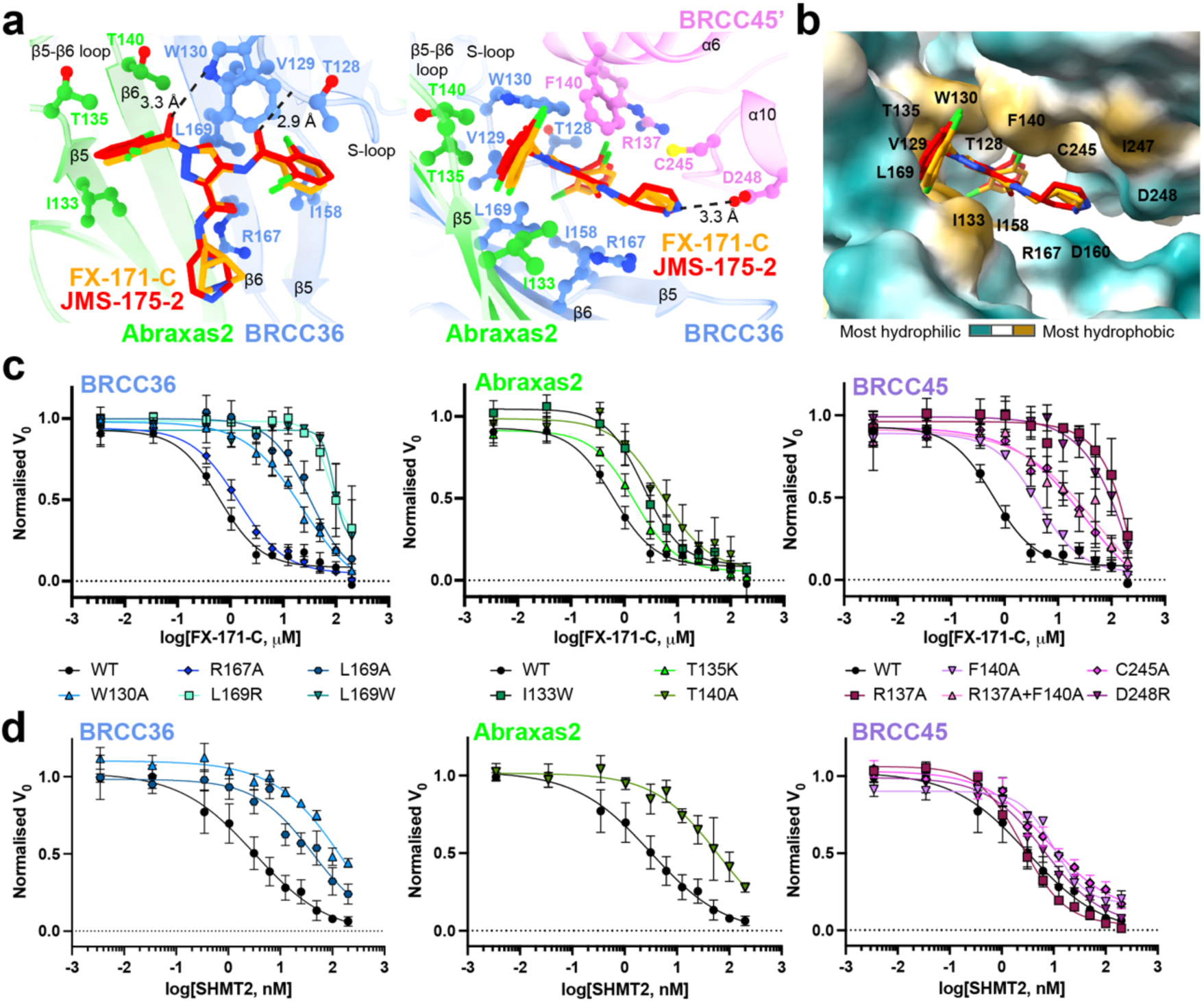
Analysis of the BLUE compound binding site. **a,** Ball and stick model of FX-171-C and JMS-175-2 binding to BRCC36, Abraxas2 and BRCC45. Hydrogen bonds shown as black dashed lines and residues studied by mutagenesis are indicated. **b,** The BLUE compound binding pocket shown as a surface and coloured by hydrophobicity. **c,** FX-171-C inhibition of BRISC DUB activity with BRCC36, Abraxas2 and BRCC45 mutants. **d**, SHMT2 inhibition of the same BRISC mutants as in **c,**. Data in **c,** and **d,** are mean ± SEM of three independent experiments carried out in technical duplicate.

BRCC36 forms two hydrogen bonds with the BLUE compounds. In both State 1 and 2, the amide backbone of V129 and the W130 sidechain (BRCC36 S-loop) form hydrogen bonds with the two amide oxygens either side of the central pyrazole ring. BRCC45’ F140 forms aromatic stacking interactions with the central pyrazole ring, and the BRCC45’ D248 forms a hydrogen bond with the amine group in the JMS-175-2 piperidine or FX-171-C pyrrolidine ring (**Fig. 3a**). Consistent with this interaction, analogues containing methyl substitutions of the piperidine ring showed reduced inhibition of BRISC activity (**Extended Data Figs. 6a, 6b**). BRCC45’ R137 forms a hydrogen bond with the BRCC45’ loop containing C245 to stabilise the BRCC45’ *α*10 helix that lines the compound binding site.

Interestingly, we do not observe the same compound binding pocket in the asymmetric (no inhibitor) conformation (**Fig. 2b**). We rigid-body fitted two BRISC molecules into the cryo-EM density of the asymmetric dimer conformation and observe a shifted BRISC’ molecule relative to the BRISC-FX-171-C model (**Fig. 2b, Extended Data. Fig 4k**). The BRCC45’ *α*6 and *α*10 helices which line the compound binding site are shifted in the compound-bound conformation (**Extended Data Fig. 4l**). These structural insights suggest that molecular glue compound binding not only induces BRISC dimerisation but also alters the conformation of pre-existing (and low-level) BRISC dimers.

**Figure 4.**
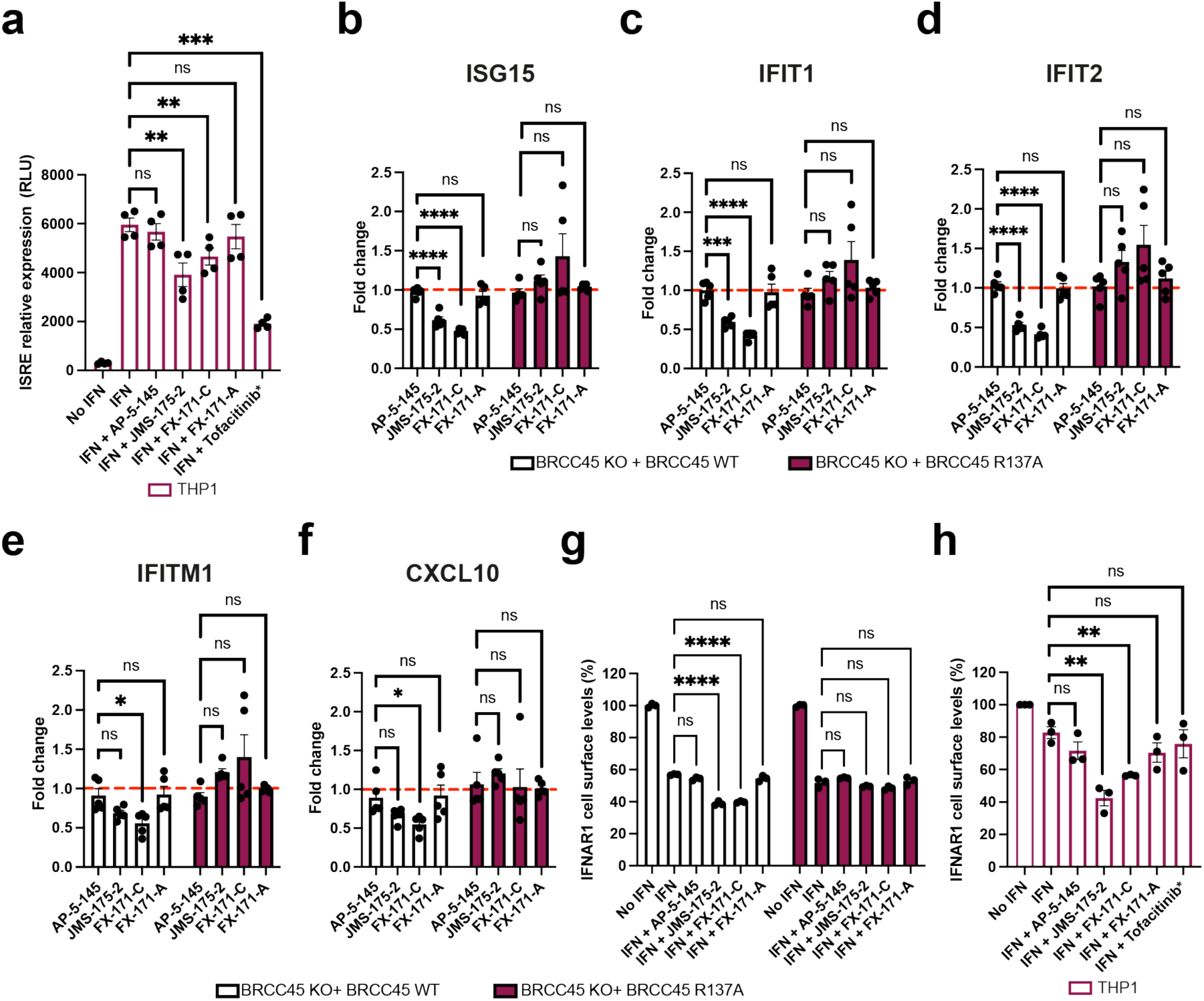
BLUE compounds reduce interferon-stimulated gene (ISG) expression and IFNAR1 internalisation in cells. **a,** THP-1 cells were treated with/without hIFNα2 (25 ng/mL) and either 4 μM inhibitor (JMS-175-2, FX-171-C, FX-171-A), 4 μM negative control AP-5-145, DMSO control (0.1%), or JAK/STAT inhibitor Tofacitinib (*0.4 μM) for 16 hours. Luciferase analysis of the ISRE in THP-1 supernatant in relative light units (RLU). Data points are from four independent experiments. **b-f,** MCF10A *Cas*9 cells expressing BRCC45 wild-type (WT) and BRCC45 R137A were treated with hIFNα2 (75 ng/mL) and either 2.5 μM inhibitor (JMS-175-2, FX-171-C, FX-171-A), 2.5 μM negative control AP-5-145 or DMSO (0.1%) for 4 hours. Expression of indicated interferon-induced genes (**b,** ISG15, **c,** IFIT1, **d,** IFIT2, **e,** IFITM1, **f,** CXCL10) normalised to 18s rRNA are presented as fold change to own IFN + DMSO treated control. Data points are from four independent experiments. **g,** MCF10A cells (BRCC45 WT and BRCC45 R137A) were treated with/without hIFN-Iα (50 ng/mL) and either 5 μM inhibitor (JMS-175-2, FX-171-C, FX-171-A), 5 μM negative control AP-5-145 or DMSO (0.1%) for 90 minutes. IFNAR1 cell surface levels (%) were quantified using FACS analysis and calculated as a percentage of no IFN stimulation. Data points are from three independent experiments. **h,** THP-1 cells were treated with/without hIFNα2 (25 ng/mL) and either 4 μM inhibitor (JMS-175-2, FX-171-C, FX-171-A), 4 μM negative control AP-5-145, DMSO control (0.1%), or JAK/SAT inhibitor Tofacitinib (*0.4 μM) for 16 hours. IFNAR1 surface levels were quantified using FACS analysis and median fluorescence intensity of allophycocyanine(APC)-IFNAR1 calculated as a percentage to no IFN stimulation. Data points are from three independent experiments. Statistical analyses in **a,** were performed using paired t-tests to compare compound treated cells to DMSO control cells. In **b-g,** one-way ANOVA with Dunnett’s multiple comparisons test was used to compare statistical significance between AP-5-145 and BLUE compound treatment. In **h,** unpaired t-tests were used to compare compound treated cells to DMSO control cells. P values illustrated by * <0.05, ** <0.01, *** <0.005, **** <0.0001, ns = non-significant. Error bars represent ± SEM.

### A human-specific BRCC36 loop promotes BRISC dimer formation

BLUE compounds are highly selective for BRISC over other JAMM/MPN DUBs, including the closely related ARISC complex which shares the BRCC36 catalytic subunit (**Figs. 1e, 1g**). Cryo-EM structures revealed BLUEs directly engage the Abraxas2 subunit. Sequence alignment of Abraxas1 (ARISC) and Abraxas2 (BRISC) illustrates divergence in the primary amino acid sequence near the BLUE compound binding site (*β*5-*β*6 loop) (**Extended Data Fig. 6c**), which likely contributes to the selectivity of BLUEs for BRISC over ARISC.

To further probe compound selectivity, we tested FX-171-C inhibition of BRISC complexes from metazoan orthologues: mouse (*Mus musculus)*, zebrafish (*Danio rerio)*, and ant (*Camponotus floridanus)*. FX-171-C has the highest potency towards human BRISC over mouse and zebrafish BRISC, whilst there is no inhibition of ant BRISC (**Extended Data Fig. 6d**). Analysing the BRISC-BLUE interaction interface explains the high specificity of BLUE compounds for human BRISC over ant BRISC. BRCC36 W130 and L169 and the Abraxas2 *β*5-*β*6 strands, which line the inhibitor binding pocket (**Fig. 3a**), are not conserved in *Cf*BRCC36 and *Cf*Abraxas2 (**Extended Data Figs. 6c, 6e**). BRCC45 C245, which contributes to compound binding, is also not conserved in *Cf*BRISC (**Extended Data Fig. 6f**).

Analysing the selectivity between human and mouse BRISC suggested a possible contributor of dimer formation. Human and mouse BRISC share over 97% sequence identity, yet FX-171-C is approximately ten times more potent as an inhibitor of human BRISC. The major difference between the two species is an extended loop region of 25 amino acids in human BRCC36 (residues 184-208) (**Extended Data Fig. 6e**). Deletion of this loop in human BRISC reduced inhibitor sensitivity by approximately 10-fold, with a similar IC_50_ for mouse BRISC which lacks the same loop (**Extended Data Fig. 6d**). Consistent with the idea that the loop region mediates dimer formation we observed fewer dimers for the human BRISC*Δ*Loop construct upon inhibitor addition in mass photometry and negative stain EM (**Extended Data Figs. 6g, 6h**). The cryo-EM density which corresponds to the BRCC36 loop extends towards an Abraxas2’ subunit from the opposing BRISC molecule, further supporting the role for this loop in mediating BRISC dimerisation (**Extended Data Fig. 6i**). Interestingly, humans have two BRCC36 isoforms with one lacking this loop region, suggesting that it is possible to design molecular glue compounds that display not only selectivity within the same enzyme family, but also across orthologous species and splice variants.

### BRISC-inhibitor interacting residues are important for inhibitor sensitivity *in vitro*

The cryo-EM structures of BRISC in complex with molecular glues allowed us to make selective mutations to probe the BRISC-BLUE interaction site and to assess the contribution of each interacting residue for inhibition. We mutated residues from BRCC36, Abraxas2, and BRCC45, and purified 15 mutant BRISC complexes from insect cells (**Extended Data Fig. 7a**). Due to the proximity of some residues to the BRCC36 active site, we assessed BRISC DUB activity against a fluorogenic di-ubiquitin substrate. Two BRCC36 mutants (T128P and I158K) were inactive, and we determined which mutations conferred a reduction in inhibitor sensitivity for the remaining 13 active mutant complexes (**Extended Data Fig. 7b**).

BRCC36 W130A and L169A/R/W mutants showed severely reduced inhibition by FX-171-C (>100 fold over WT complex), whilst BRCC36 R167A remained inhibitor sensitive (**Fig. 3c, Extended Data Fig. 7c**). Abraxas2 mutant T140A was moderately affected, exhibiting an IC_50_ 10-fold higher than BRISC WT, while Abraxas2 I133W and T135K had moderate to little effect on inhibitor sensitivity (**Fig. 3c, Extended Data Fig. 7c**). We also mutated BRCC45’ residues (R137A, F140A, C245A, D248R) and all had reduced sensitivity to inhibition when compared with WT BRISC complexes (**Fig. 3c, Extended Data Fig. 7c**). These data validate the BLUE compound binding sites identified by cryo-EM, and the reduced sensitivity observed for the BRCC45’ mutants confirms the molecular gluing mechanism of inhibition.

The reduced inhibition for residues surrounding the inhibitor binding pocket is consistent with regions of change in solvent accessibility observed by HDX-MS after incubation with FX-171-C. BRCC36 W130 is encompassed within a protected loop (residues 111-135), and the BRCC36 W130A mutant has reduced inhibitor sensitivity (**Fig. 3c, Extended Data Fig. 7d**). Mutations of BRCC45 residues R137, F140, C245, and D248 lead to reduced FX-171-C inhibition, and these residues are in close proximity to protected regions of BRCC45: 122-134 and 206-221 (**Fig. 3c, Extended Data Fig. 7d**). In contrast, BRCC36 L169 and Abraxas2 T140 are required for inhibition but do not exhibit changes in solvent accessibility in HDX-MS.

### Molecular glues and SHMT2 share a binding pocket

The metabolic enzyme SHMT2 interacts with BRISC to regulate IFNAR1 signalling^20^. Interestingly, the SHMT2 binding site on BRISC overlaps with the BLUE compound binding site (**Extended Data Fig. 7e**). Indeed, some of the residues we mutated to validate inhibitor binding also contribute to the SHMT2 interaction interface. As SHMT2 is a potent endogenous inhibitor of BRISC DUB activity^20^, we assessed if BRISC mutants were still inhibited by SHMT2 (**Fig. 3d**). BRCC36 W130A and L169A and Abraxas2 T140A show reduced BRISC inhibition by SHMT2, indicating these mutations also disrupt SHMT2 binding to BRISC. By contrast, the BRCC45 mutants were inhibited by SHMT2 with a similar IC_50_ to BRISC WT, which is consistent with these BRCC45 residues being far away from the BRISC-SHMT2 binding interface in the context of the BRISC monomer (**Extended Data Fig. 7f**). Therefore, both the BLUE compounds and the endogenous inhibitor SHMT2 share a common interacting site.

To investigate SHMT2 and BLUE compound competition with BRISC, we used a Spectral Shift assay (Dianthus) to determine the *K*_D_ of BRISC-SHMT2 interaction in the absence and presence of FX-171-C. We measured a *K*_D_ of 0.4 ± 0.1 μM for SHMT2 with labelled BRISC (**Extended Data Fig. 7g**). The affinity of SHMT2 for BRISC was reduced to 3.6 ± 1.8 μM after incubation with FX-171-C, and 0.7 ± 0.3 μM with JMS-175-2. There was no change in affinity with the negative control compound AP-5-144 (**Extended Data Fig. 7g**). These data demonstrate direct competition between BLUEs and SHMT2 and show BLUEs can reduce SHMT2 binding to BRISC which is required for immune signalling in cells^20^.

### BLUE compounds reduce interferon signalling

To investigate the effects of BLUE compounds on interferon signalling we used the THP-1 cell line, which contains a stably integrated, inducible luciferase reporter construct for the interferon regulatory factor (IRF) pathway. For additional controls alongside FX-171-C and JMS-175-2, we tested three further compounds: FX-171-A, AP-5-145 and Tofacitinib, a JAK inhibitor. FX-171-A has a similar chemical structure to JMS-175-2 but is a less potent BRISC inhibitor (IC_50_ = 6.8 μM) (**Extended Data Figs. 8a, 8b**). AP-5-145 is an N-methylated analogue of AP-5-144, the other possible stereoisomer synthesised at the beginning of this study and has no inhibitory effect against BRISC (**Fig. 1c, Extended Data Fig. 8b**). We added a methyl group to the pyrazole ring to avoid potential CDK2 inhibition in cells and to increase cell permeability when used as a negative control. We observed no cell death with any of the compounds tested up to 4 μM (**Extended Data Fig. 8c**).

We first evaluated the impact of BLUE compound treatment on the activation of interferon-stimulated response elements (ISRE) in response to IFNα2 stimulation. Both JMS-175-2 and FX-171-C reduced ISRE relative expression compared to the AP-5-145 and FX-171-A control compounds, with Tofacitinib having a potent effect (**Fig. 4a**).

Next, we stimulated THP-1 cells with different agonists to determine if BLUE compound treatment affected other signalling pathways. We stimulated THP-1 cells with TLR3 and TLR9 agonists, polyinosinic:polycytidylic acid (poly I:C) and ODN 2216. These only minimally induced ISRE relative expression, which was not reduced by BLUE compound or Tofacitinib treatment (**Extended Data Figs. 8d, 8e**). In addition, we used a NF-κB pathway reporter assay and observed no BLUE compound effect on LPS-induced NF-κB pathway activity compared to the control compounds (**Extended Data Fig. 8f**).

To probe the on-target effect of BLUE compounds in cells we generated BRCC45 KO MCF10A cells using CRISPR-*Cas*9-mediated genomic deletion (**Extended Data Fig. 8h**) and complemented BRCC45 KO with either Flag-BRCC45 WT or Flag-BRCC45 R137A (**Extended Data Fig. 8i**). In DUB activity assays, BRCC45 R137A mutation reduced FX-171-C inhibition (IC_50_ > 100 µM) without affecting SHMT2 inhibition (**Figs. 3c, 3d, Extended Data Fig. 7c**). We confirmed by co-immunoprecipitation that interactions with BRISC subunits BRCC36 and MERIT40 were maintained in the BRCC45 WT and BRCC45 R137A cell lines (**Extended Data Fig. 8j**).

MCF10A cells were challenged with IFNα2 to stimulate IFNAR1 signalling and ISG expression. An increase in STAT1 phosphorylation was observed in the sgROSA (control sgRNA) cell line, BRCC45 WT, and BRCC45 R137A cell lines, and less STAT1 phosphorylation was observed in BRCC45 KO cells (**Extended Data Fig. 8k**). Following IFNα2 stimulation, we used quantitative PCR with reverse transcription (qRT-PCR) to measure changes in gene expression for five ISGs: ISG15, IFIT1, IFIT2, IFITM1, and CXCL10. The differences in gene expression between sgROSA, WT and R137A cell lines were non-significant, except for ISG15, which had higher gene expression in BRCC45 WT cells, but still comparable to BRCC45 R137A (**Extended Data Fig. 8l**).

To measure the effect of BLUE compounds on ISG expression, we compared BRCC45 WT and BRCC45 R137A cell lines with active and inactive control compounds. Treatment of BRCC45 WT cells with 2.5 μM JMS-175-2 and FX-171-C reduced gene expression for all five ISGs (**Figs. 4b-f**). Critically, a reduction in ISG expression was not observed in cells harbouring the BRCC45 R137A mutation, which reduces BLUE inhibition of BRISC, indicating on-target BLUE activity in cells.

BRISC regulates interferon signalling through IFNAR1 deubiquitylation, likely limiting IFNAR1 internalisation and subsequent degradation^23^ (**Extended Data Fig. 10**). We used fluorescence-activated cell sorting (FACS) to determine the effect of BLUE compounds on IFNAR1 surface levels. MCF10A cell lines were challenged with IFNα2, and reduced IFNAR1 surface levels were observed (**Extended Data Fig. 8m**), indicating increased IFNAR1 internalisation in response to IFN stimulation. We treated the sgROSA and BRCC45 WT cell lines with the compound panel, and observed a reduction in IFNAR1 cell surface levels with FX-171-C and JMS-175-2, but not with FX-171-A and AP-5-145 (**Fig. 4g, Extended Data Fig. 8g**). Importantly, IFNAR1 surface levels were not reduced in the BRCC45 R137A cells, indicating reduced IFNAR1 levels in sgROSA and BRCC45 WT cells are due to BRISC inhibition (**Fig. 4g**). Similarly, we stimulated the monocyte cell line THP-1 with IFNα2 and treated with the inhibitor panel and analysed IFNAR1 levels using FACS. We observed a reduction in IFNAR1 surface levels after treatment with JMS-175-2 and FX-171-C and no significant reduction was observed with AP-5-145 or FX-171-A, or JAK/STAT inhibitor Tofacitinib (**Fig. 4h**).

To evaluate the impact of BLUE compound treatment and BRISC inhibition on IFNAR1 ubiquitylation, we used tandem ubiquitin binding entities (TUBE) and Western blotting. Following stimulation with IFNα2 in MCF10A cells, we isolated ubiquitylated IFNAR1 via TUBE pull-downs. We observed elevated levels of IFNAR1 ubiquitylation with FX-171-C and JMS-175-2 treatments compared to treatment with DMSO, AP-5-145, and FX-171-A (**Extended Data Fig. 8n**).

We next studied the effects of BLUE compound treatment in peripheral blood mononuclear cells (PBMCs) from healthy volunteers upon IFNα stimulation. For a comprehensive analysis of how BLUE compounds modulate the IFN gene signature, we measured the expression of 67 ISGs, including ISG15, IFIT1, IFIT2, IFITM1, CXCL10 and MX1. 34 ISGs were significantly upregulated by IFN, with no difference in ISG expression profile between DMSO or AP-5-145 (**Fig. 5a, Extended Data Figs. 9c, 9d**). Treatment with JMS-175-2 suppressed the IFN-induced signature compared to AP-5-145 in 13 of the 34 ISGs (**Fig. 5b, Extended Data Fig. 9a**). Interestingly, for FX-171-C-treated PBMCs, there was less reduction in ISG expression compared to AP-5-145 control when used at 2 μM (**Extended Data Fig. 9a**). We also measured IFN-inducible CXCL10 secretion levels and observed lower CXCL10 protein secretion in BLUE-treated PBMCs (**Fig. 5c**).

**Figure 5.**
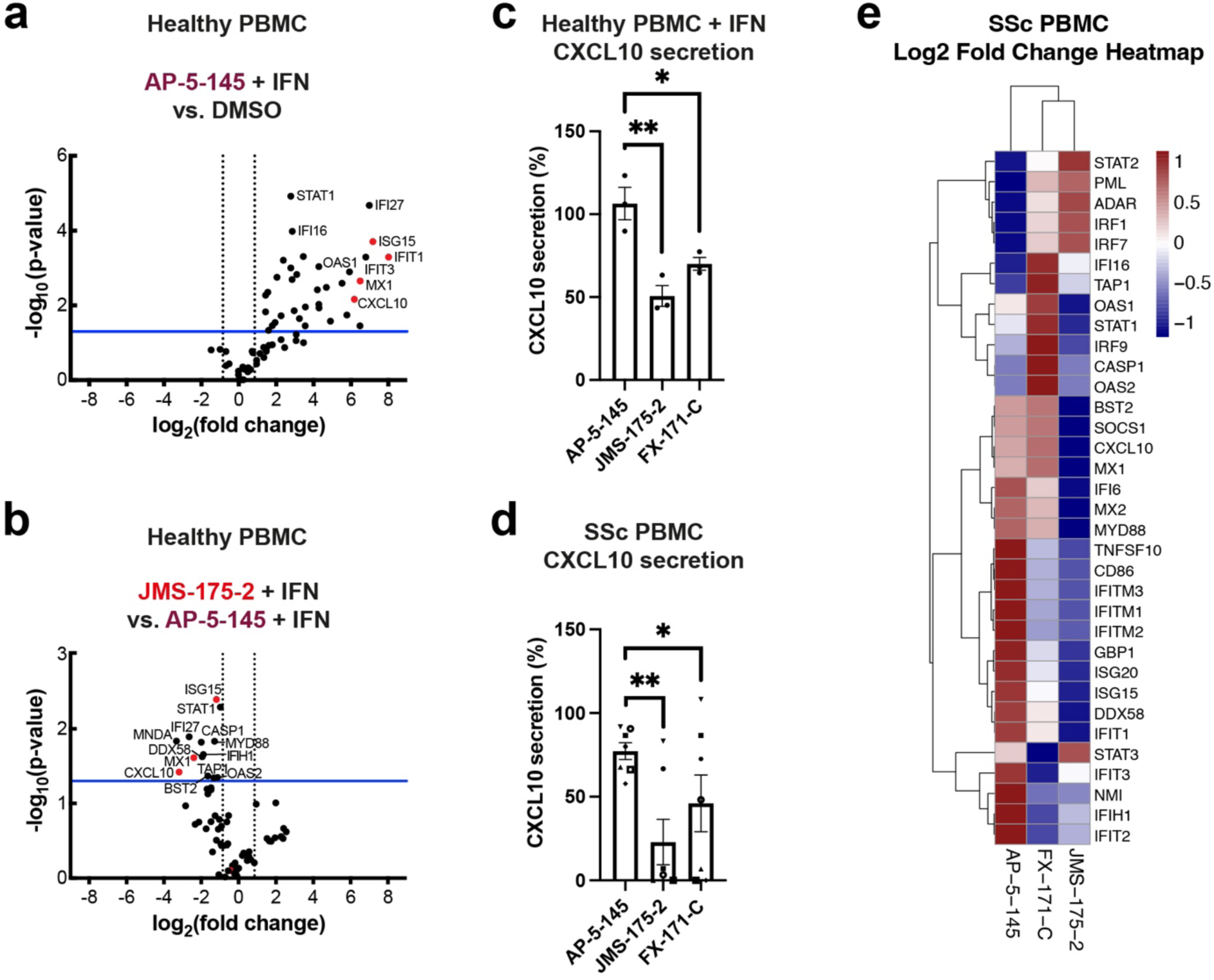
BLUE compounds reduce interferon-stimulated gene (ISG) expression in PBMCs. **a, b,** Type I IFN signalling gene expression analysis (67 genes normalised for housekeeping genes: ACTB, GAPDH, HPRT1, RPLP0) of healthy control PBMCs stimulated with IFNα2. Volcano plot of genes increased with addition of IFNα2 with negative control AP-5-145 compared to DMSO (no IFN) condition, and **b,** effect of JMS-175-2 + IFN stimulation compared to AP-5-145 + IFN. Blue line indicates a p-value of 0.05. Data points are the means from three independent experiments. **c,** CXCL10 protein levels in supernatant from IFNα2-stimulated healthy PBMCs (n=3) quantified by ELISA and shown as a percentage of own IFN + DMSO control (100%). Bar graph is average of three independent experiments. **d,** PBMCs were isolated from SSc patients and treated with DMSO, AP-5-145, FX-171-C, or JMS-175-2 for 16 hours without IFN stimulation. Secreted CXCL10 in supernatant is shown as percentage to own DMSO control. Bar graph is average of data from seven SSc donors. **e,** Type I IFN signalling gene expression analysis of unstimulated SSc PBMCs, treated with 2 µM AP-5-145, JMS-175-2, or FX-171-C. Heat map showing log2 mean fold change in ISG expression by treatment, compared to AP-5-145, via qPCR SuperArray. ΔCt was calculated against the geometric mean of four housekeeping genes, followed by ΔΔCt (fold change) relative to AP-5-145, and log2 transformation. Heat maps represent the mean fold change from nine SSc donors. In **a,** and **b,** paired two-tailed student t-tests were used to compare between treatment conditions for statistical significance. P values illustrated by * <0.05, ** <0.01, ns = non-significant. Error bars represent ± SEM.

To assess the activity of BLUE compounds in the context of autoimmune disease associated with aberrant Type I IFN activation, we also measured ISG expression and CXCL10 secretion levels of PBMCs from patients affected by Scleroderma (SSc)^26^ (**Extended Data Table 3**). Treatment of basal, unstimulated SSc PBMCs with JMS-175-2 or FX-171-C reduced CXCL10 secretion (**Fig. 5d**). Next, we used qRT-PCR to assess the effect of BLUE compounds to reduce Type I IFN signalling in non-stimulated SSc PBMCs, by analysing the expression of the 34 ISGs determined in healthy PBMCs (**Fig. 4a**). In SSc PBMCs (n=9), 22 ISGs were reduced after treatment with JMS-175-2 compared to AP-5-145, with FX-171-C having a lesser effect (**Fig. 5e, Extended Data Fig. 9b**). Additionally, we used a composite ISG score including CXCL10, IFIT1, ISG15 and MX1 relative to GAPDH^48^ to measure the effect on ISG expression for a larger group of patients (n=20). BLUE treatment of unstimulated SSc PBMCs reduced the composite ISG score for JMS-175-2 and FX-171-C relative to AP-5-145 (**Extended Data Fig. 9e**).

## Discussion

Deubiquitylating enzymes are attractive drug targets due to their roles in cancer, neurodegeneration, inflammation, and immunity^10,49–51^. The BRISC DUB complex regulates Type I interferon signalling and is a promising target for ameliorating autoimmune disease conditions^24–26^. However, the high conservation among active sites of JAMM/MPN DUBs makes discovery of selective inhibitors challenging. Moreover, the same catalytic subunit (BRCC36) is found in two structurally related protein complexes with separate functions. Using a combination of biochemical screening, structural and molecular biology techniques, we identified first-in-class, selective inhibitors of the cytoplasmic BRISC DUB complex over structurally related DUBs.

BRISC inhibitors are molecular glues that induce the formation of an inhibited BRISC dimer. We showed target engagement in cells using an inhibitor resistant mutation, BRCC45 R137A. In addition, we measured a reduction in interferon-stimulated gene expression in MCF10A and THP-1 cells and PBMCs from healthy and scleroderma patients after treatment with BLUE compounds. While we observe a reduction of ISG levels with JMS-175-2 treatment when looking at averaged patients data (**Fig. 5e**), patient-to-patient variability masks this moderate effect in individual analyses (**Ext. Data Fig. 9b**). A key finding in cellular studies is that BLUEs inhibit the pathway by affecting interferon receptor surface levels rather than blocking IFN signalling directly, consistent with being first-in-class JAMM domain DUB inhibitors. We recognise that the magnitude of pathway inhibition is modest, especially when compared to established clinical drugs such as JAK inhibitors. While this effect is likely dependent on BLUEs potency, which could be improved in future drug development efforts, it is tempting to speculate that a partial inhibition of the pathway may be exploited for normalising rather than neutering Type I IFN activation in disease conditions.

Based on current and published reports about IFNAR1 being a substrate of BRISC^20,23^, we propose a model where reduced BRISC activity results in hyper-ubiquitylation of IFNAR1 and accelerated receptor degradation, leading to reduced inflammatory signalling (**Extended Data Fig. 10**). Combining structural analysis of current models of BRISC interactions with SHMT2 and ubiquitin, we propose that BLUEs stabilise an autoinhibited BRISC dimer.

DUBs are regulated through autoinhibited conformations, protein-protein interactions, and interactions with pseudo-DUB partners. To our knowledge this is the first example of a DUB molecular glue. This concept could be applied to other DUBs or macromolecular complexes to develop selective inhibitors by promoting interactions instead of using the classical approach of breaking protein-ligand or protein-protein interactions. Molecular glues which stabilise autoinhibited conformations offer a unique opportunity to exploit naturally existing mechanisms, for example, NX-1607, an inhibitor of the Cbl-b E3 ligase, is an intramolecular glue which traps the ligase in an inactive state^52,53^. Moreover, this approach defines a route to achieve selectivity for DUBs with highly conserved active sites and which also exist as dimers (e.g. USP25 and USP28^54,55^).

Indeed, there are several examples of USP inhibitors which block catalysis by stabilising an inactive enzyme conformation. USP7 inhibitors FT671 and FT827^28^ and the cross-reactive USP11 and USP15 inhibitor mitoxantrone^56,57^ achieve inhibition through misalignment of active site residues. Other USP7 inhibitors, GNE-6640 and GNE-6676 inhibit by blocking ubiquitin binding rather than disrupting the catalytic site^58^. Similarly, BLUEs block catalysis without disrupting the BRCC36 catalytic site (**Extended Data Fig. 5d**). This is the first example of a new class of JAMM/MPN DUB inhibitor. All other JAMM/MPN inhibitors, including the selective CSN5 inhibitors^34^, interact with the catalytic zinc (**Extended Data Fig. 5e**).

BLUE compounds achieve selectivity by exploiting a binding site at the interface of three different proteins (**Fig. 2e**). Most of the currently identified molecular glues contact two different proteins, with a few exceptions. Natural compounds Cyclosporin A (CsA) and FK506 were the first characterised molecular glues and exert immunosuppressive activity by disrupting signalling events mediated by calcineurin^38^. CsA and FK506 stabilise interactions between three different proteins. Both compounds contact calcineurin A and calcineurin B. FK506 induces an interaction with FKBP12 and CsA induces an interaction with cyclophilin^38,59,60^. Interestingly, in the case of the BLUE compounds, they interact at the interface of a DUB, a pseudo-DUB and a UEV domain, contacting a previously unexplored UEV site.

DUB complexes use protein-protein interactions to regulate enzymatic activity, cellular localisation, and substrate binding. The subcellular localisation, function and activity of BRCC36 is regulated by the interaction of pseudo-DUBs, namely Abraxas1 and Abraxas2^16^. Here, we show additional regulation of BRISC activity through dimerisation. The identification of BRISC dimers in the absence of small molecules alludes to a mechanism of action whereby the molecular glues stabilise a low affinity interaction between BRISC molecules. Indeed, the ability of molecular glues to enhance the affinity between proteins which already have a pre-existing low affinity interaction is what distinguishes them from bifunctional molecules such as PROTACs^61^. The asymmetric dimer conformation blocks the BRCC36 active site and predicted polyubiquitin binding sites (**Extended Data Fig. 3d**). Moreover, the BLUE binding pocket overlaps with the binding interface of the endogenous inhibitor SHMT2 (**Extended Data Fig. 7e**). Therefore, it is possible that the asymmetric dimer represents an autoinhibited BRISC conformation which is sampled by a small population of the complex in the absence of external stimuli (i.e. small molecule binding or as yet, unidentified mechanisms). Determining the functional importance of the different BRISC dimers and identifying their potential regulation is an important area of future exploration.

Testing BLUE inhibition of BRISC orthologues highlighted a human-specific BRCC36 loop which is important for inhibitor sensitivity and dimer formation. A human splice isoform also lacks this loop region (residues 184-208). In cryo-EM maps of the BRISC dimer, this flexible BRCC36 loop extends towards the opposite BRISC molecule and acts to stabilise the BRISC dimer (**Extended Data Fig. 6i**). These insights suggest the autoinhibited conformation is prevalent for the human BRISC complex and partially driven by the human-specific BRCC36 loop. Further investigation using potent inhibitors with better drug-like properties are required to elucidate the regulation and formation of BRISC dimers in cells and animals. The difference between mouse and human forms needs special consideration if murine models are used for efficacy or translational studies. Additionally, humanised mouse models may be necessary to explore inhibitor effects in vivo, and to elucidate the potential for BRISC inhibitors as future therapeutics.

## Materials and methods

### Reagents

Reagents and antibodies were purchased from the following commercial sources: Anti-Flag M2 Affinity Gel beads (Sigma-Aldrich, #A2220), Recombinant Human IFNα2 (carrier free) (Biolegend, #592702) were purchased for MCF10A experiments. The following antibodies were used for immunoblotting: Anti-BRCC45 (Abcam, #ab177960), Anti-GAPDH (Cell Signaling Technology, #2118S), Anti-Merit40 (Cell Signaling Technology, #12711S), Anti-BRCC36 (Abcam, #ab108411), Anti-Phospho-Stat1 (Tyr701) (Cell Signaling Technology, #9167S), Anti-α Actin (Santa Cruz Biotechnology, SC-#32251), anti-IFNAR1 antibody (ab124764, Abcam). POWER RT-qPCR reagents including SYBR Green PCR Master mix (#4367659) and cDNA Reverse Transcription Kit (#4368814) were purchased from Allied Biosystems. RNeasy mini kit (#74104) and RNase free DNase (#79254) were purchased from Qiagen. The 4–12% Bis-Tris SDS-PAGE gels, cell culture media, horse serum and other materials for tissue culture were purchased from Invitrogen. LipoD293 (SigmaGen, #SL100668) was used for all the transfections to generate knock out cell lines. Media supplements including human insulin solution (Santa Cruz Biotechnology, #sc-360248), cholera toxin from vibrio (Sigma Aldrich, #C8052-2MG), recombinant human EGF (PeproTech, #AF-100-15-100μg), hydrocortisone (Sigma-Aldrich, #H-0888) were used to grow MCF10A cells.

### Expression and purification of DUB complexes

Four-subunit human BRISC and ARISC complexes (BRISC full-length (FL), ARISC (FL), BRISC*Δ*N*Δ*C (MERIT40*Δ*N, Abraxas2*Δ*C), BRCC36-Abraxas2, BRISC*Δ*Loop(*Δ*184-208), *Dr*BRISC*Δ*N*Δ*C, *Cf*BRISC*Δ*N*Δ*C) were cloned using the MultiBac system and co-expressed in *Spodoptera frugiperda* (*Sf*9) insect cells^62^. BRISC mutants and *Mm*BRISC were cloned into pFastBac-HTB vectors in the Bac-to-Bac system (ThermoFisher Scientific), baculoviruses were generated in *Sf*9 cells and used for co-infection of *Trichoplusia ni* (*Tni*) cells. All BRISC complexes were purified as previously described^18,20^.

USP2 with an N-terminal His-tag was purchased from Addgene (plasmid #36894). AMSH*, an AMSH-STAM fusion^42^, with an N-terminal His-tag was purchased from Addgene (pOPINB-AMSH*, plasmid #66712). SHMT2*Δ*N(A285T) (residues 18-504) was expressed and purified as previously described^20^. His-USP2 and His-AMSH* were expressed in *Escherichia coli* BL21 (DE3) cells. Cells were grown at 37 °C in Terrific Broth (TB) medium, induced with 0.5 mM isopropyl-β-D-1-thiogalactopyranoside (IPTG) and grown overnight at 18 °C. For purification, cell pellets were resuspended in lysis buffer containing 50 mM Tris-HCl pH 7.6, 300 mM NaCl, 20 mM imidazole, 5% glycerol, 0.075% β-mercaptoethanol, 1 mM benzamidine, 0.8 mM phenylmethylsulfonyl fluoride (PMSF) and 0.3 mg/mL lysozyme. Cells were lysed by sonication (1 s on, 1 s off for a total of 16 minutes) and cleared by centrifugation at 18,000 g. The clarified lysate was incubated with Ni-NTA beads (Cytiva) for 1 hour at 4 °C, before washing with wash buffer containing 50 mM Tris-HCl pH 7.6, 300 mM NaCl, 20 mM imidazole, 5% glycerol, 0.075% β-mercaptoethanol, and 1 mM benzamidine and a high salt buffer containing 500 mM NaCl. The protein was eluted with elution buffer containing 50 mM Tris-HCl pH 7.6, 300 mM NaCl, 120 mM imidazole, 5% glycerol, 0.075% β-mercaptoethanol, and 1 mM benzamidine. The elutions containing His-USP2 or His-AMSH* were dialysed overnight with thrombin (His-USP2) or 3C PreScission protease (His-AMSH*). After dialysis, cleaved samples were incubated with Ni-NTA beads and washed with wash buffer. The cleaved fractions were concentrated and loaded onto a Superdex 75 10/300 column (Cytiva) equilibrated with 25 mM HEPES pH 7.5, 150 mM NaCl, and 1 mM tris(2-carboxyethyl)phosphine (TCEP).

### Compound library screening

BRISC DUB activity was measured at room temperature (RT) using 1 nM BRISC and 500 nM internally quenched fluorescence (IQF) diUb K63 substrate (Lifesensors, DU6303) in the presence of DMSO and compounds at 10 μM (final concentrations). Assays were performed in 384-well black flat-bottom low-flange plates (Corning; 35373) in buffer containing 50 mM HEPES-NaOH pH 7.0, 100 mM NaCl, 1 mg/mL bovine serum albumin (BSA), 1 mM dithiothreitol (DTT), and 0.03% v/v Brij-35. Ten microlitres of x2 concentrated enzyme stock was dispensed followed by transfer of 200 nL of compounds (1 mM stock) using a 384-pin tool and a 15 min incubation at RT. Ten microlitres of x2 concentrated substrate stock was added and the reaction was monitored by measuring fluorescence intensity (excitation, 540 nm; emission, 580 nm) after 20 min incubation at RT and ∼50% of substrate was consumed. Orthogonal assays for verification of H20 and P12 hit compounds using USP2 (100 nM) and Trypsin (125 nM) were performed in identical conditions.

### LCMS and MS-MS analysis

LC MS-MS analysis was carried out in a UPLC BEH C18 (2.1 × 50 mm, 1.7 µm) column using ACQUITY UPLC II system. The mobile phase was 0.1% formic acid in water (solvent A) and 0.1% formic acid in acetonitrile (solvent B). A gradient starting at 95% solvent A going to 5% in 4.5 minutes, holding for 0.5 minutes, going back to 95% in 0.5 minutes and equilibrating the column for 1 minute was employed. A Waters Synapt G2S QTof mass spectrometer equipped with an electrospray ionization source was used for mass spectrometric analysis. MassLynx (v4.1) was used for data analysis. The MS parameters for LCMS analysis were frequency of 15 s, cone voltage of 25 V, capillary voltage was 3 kV. For MS-MS spectra, an MS-MS range of m/z 50-900, scan time of 0.1 s, collision energy ramp of 30-60 volts were used. Compound identifications were performed using accurate mass analysis and MS-MS fragmentation analysis. Absorbance spectra was measured using a UV-VIS nanodrop spectrophotometer N8000 (ThermoFisher Scientific).

### Synthesis of BRISC inhibitors

Reaction schemes, methods, and validation of the synthesis of JMS-175-2, FX-171-C, AP-5-145, and FX-171-A are outlined in the **Supplementary Information**.

### DUB activity assays (IQF)

IQF assays were performed in DUB reaction buffer containing 50 mM HEPES-NaOH pH 7.0, 100 mM NaCl, 0.1 mg/mL BSA, 1 mM DTT, and 0.03% v/v Brij-35. To assess inhibitor potency, inhibitors were diluted in DMSO up to a final concentration of 200 μM and incubated with the target enzyme for 15 minutes at RT. The final concentration of BRISC WT, all BRISC mutants, *Hs*BRISC*Δ*Loop, *Mm*BRISC and *Dr*BRISC*Δ*N*Δ*C was 1 nM. Other enzymes were tested at the following concentrations: 5 nM ARISC (FL), 10 nM *Cf*BRISC*Δ*N*Δ*C, 250 nM AMSH* (a STAM2-AMSH fusion^42^), 500 nM USP2, and 1 μM trypsin (Sigma, 9002-07-7).

Michaelis-Menten analysis in the absence and presence of increasing concentrations of either JMS-175-2 or FX-171-C were performed in DUB reaction buffer. BRISC concentration was 1 nM and IQF K63-linked di-ubiquitin substrate was used from 0 to 1 μM. Michaelis-Menten analyses and Lineweaver-Burk plots were calculated using GraphPad Prism (v9.0).

To assess enzymatic activity, BRISC complexes were diluted to concentrations between 1 nM and 50 nM. For SHMT2 assays, SHMT2(A285T) was diluted in 20 mM MES pH 6.5, 500 mM NaCl and 2 mM TCEP up to a final concentration of 200 nM. SHMT2 was incubated with enzyme for 15 minutes at RT before adding substrate. 20

μL enzyme reactions were carried out in 384-well black flat-bottom low flange plates (Corning; 35373). DUB activity was measured using internally quenched fluorescent (IQF) K63-linked diUb (Lifesensors, DU6303) at 50 nM. Cleaved di-ubiquitin was monitored by measuring fluorescence intensity (Ex. 544 nm, Em. 575 nm; dichroic mirror 560 nm). Fluorescence intensity was measured every minute for 30 minutes at 30 °C. Fluorescence intensity units were plotted against time to generate a linear reaction progress curve, where the initial velocity (V_0_) corresponds to the gradient of the curve (typically achieved within 15 minutes). IC_50_ values were calculated using the GraphPad Prism (v9.0) built-in dose-response equation for inhibitor concentration vs. response (variable slope).

### DUB selectivity profiling

Biochemical selectivity profiling (DUBprofiler^TM^) was performed by Ubiquigent with a panel of 48 purified DUBs and ubiquitin-rhodamine(110)-glycine as a fluorescent substrate. Single-dose inhibition (5 μM) was determined for JMS-175-2 after 15 minutes pre-incubation at RT.

### Deubiquitylation assay using fluorescently labelled polyUb chains

The inhibitor capzimin (Merck) was diluted in 1 mM DTT and incubated for 30 minutes at RT to reduce disulphide bonds required for inhibitor activity. 5 nM BRISC (FL) or 20 nM ARISC (FL) were incubated with either 0.5% DMSO, or 10 μM or 100 μM inhibitor (FX-171-C, JMS-175-2, AP-5-144, thiolutin or capzimin) for 15 minutes at RT in DUB reaction buffer containing 50 mM HEPES-NaOH pH 7.0, 100 mM NaCl, 0.1 mg/mL BSA, 1 mM DTT, and 0.03% v/v Brij-35. DUBs were incubated with 750 nM TAMRA-labelled K63-tetraubiquitin (LifeSensors; SI6304T) for 10 minutes at 30 °C. Reactions were quenched by adding 2.5 μL 4X SDS-PAGE loading dye (240 mM Tris-HCl pH 6.8, 40% v/v glycerol, 8% w/v SDS, 0.04% w/v bromophenol blue, 5% v/v *β*-mercaptoethanol). The samples were resolved on 4-12% NuPAGE Bis-Tris gels (ThermoFisher Scientific). Gels were scanned using an iBright FL 1500 (ThermoFisher Scientific) (Ex. 515-545 nm, Em. 568-617 nm, 500 ms exposure).

### Mass photometry

For single-point measurement of BRISC and inhibitors, 1 μM BRISC(FL) was mixed with either DMSO, JMS-175-2, or FX-171-C at 330 μM and incubated for 15 minutes on ice. Immediately prior to mass photometry (MP) measurement, the BRISC-inhibitor mix was diluted in 25 mM HEPES pH 7.5, 150 mM NaCl, 1 mM TCEP to a final concentration of 10 nM BRISC (0.05% v/v DMSO). 12 μL of the diluted sample were used for the final MP measurement, following autofocus stabilisation. For measurements with increasing concentrations of JMS-175-2 and FX-171-C, 2-fold dilutions of inhibitor in 100% v/v DMSO generated a dilution series with concentrations of inhibitor from 800 μM to 0 μM. 0.5 μL inhibitor was mixed with 19.5 μL 50 nM BRISC*Δ*N*Δ*C (2.5% v/v DMSO) and incubated at RT for 15 minutes. The BRISC-inhibitor mix was used directly for MP measurement using buffer-free autofocus stabilisation.

Microscope coverslips were prepared as previously described^63^. All mass photometry experiments were performed using a One^MP^ mass photometer (Refeyn). Movies were recorded for 60 seconds using AcquireMP (Refeyn), and were processed using DiscoverMP (Refeyn). Mass photometry image processing has been previously described^63^. Briefly, contrast-to-mass (C2M) calibration was performed using protein standards (66-669 kDa) diluted in gel filtration buffer (25 mM HEPES pH 7.5, 150 mM NaCl, 1 mM TCEP). The output from each individual movie resulted in a list of particle contrasts which were converted to mass using the C2M calibration. The mass distribution from each run is in a histogram, where count refers to each landing event and a Gaussian sum is fitted to the data. The relative amount of each species is calculated as the area of each Gaussian, where σ refers to the standard deviation of the fitted Gaussian. Dimer fraction refers to the percentage of the total counts which correspond to the 564 kDa BRISC*Δ*N*Δ*C dimer complex. Curves and EC_50_ values were fitted and calculated using the GraphPad Prism (v9.0) built-in dose-response equation for concentration of agonist vs. response (variable slope).

### Negative stain electron microscopy – grid preparation, data collection and image processing

BRISC(FL) or *Mm*BRISC(FL) was mixed with inhibitor (or DMSO) on ice for 30 minutes at a final concentration of 1 μM BRISC and 100 μM inhibitor. The BRISC-inhibitor mix was diluted in gel filtration buffer to a final concentration of 0.014 mg/mL (0.5% v/v DMSO). Sample was immediately loaded onto carbon-coated copper grids (Formvar/Carbon 300 mesh Cu, Agar Scientific). Grids were glow discharged for 30 seconds, at 10 mA, and 0.39 mBar pressure (PELCO easiGlow, Ted Pella). Grids were incubated for 1 minute with 7 μL sample, washed three times with H_2_O, stained twice with 2% w/v uranyl acetate for a total of 30 seconds. Excess liquid was removed by blotting with filter paper. Data were collected using an FEI Tecnai F20 microscope (ThermoFisher) at 200 KeV, fitted with an FEI CETA (CMOS CCD) camera. Micrographs were collected at 29000x magnification with a pixel size of 3.51 Å. RELION (v3.0 and v3.1) were used for processing of negative stain EM data^64,65^. Approximately 2,000 particles were manually picked and extracted with a box size of 128 Å^2^. These particles were used for reference-free 2D class averaging to generate 2D templates for autopicking. The parameters for autopicking were optimised and between 5,000-10,000 particles were extracted. Two rounds of 2D classification were used to remove junk particles and assess the stoichiometry of the BRISC complex.

### Cryo-electron microscopy grid preparation and data collection

To obtain the cryo-EM maps of the BRISC(FL) monomer and dimer in absence of compound, Quantifoil R1.2/1.3 300 mesh copper grids were glow discharged using a PELCO easiGlow glow discharge system (TedPella). 3 μL BRISC(FL) at 0.7 mg/mL in gel filtration buffer was loaded onto the cryo-EM grids. For the BRISC-JMS-175-2 grid, BRISC*Δ*N*Δ*C at 0.3 mg/mL (2 μM) was mixed with JMS-175-2 at 200 μM in gel filtration buffer for 30 minutes on ice. Quantifoil R1.2/1.3 300 mesh copper grids were glow-discharged using a GloQube (Quorum) for 30 seconds at 40 mA. For the BRISC-FX-171-C grid, BRISC*Δ*N*Δ*C at 0.7 mg/mL (5 μM) was mixed with FX-171-C at 400 μM in gel filtration buffer, and loaded onto grids which were plasma cleaned in downstream mode at radio-frequency power 43 W for 30 seconds using a Tergeo plasma cleaner (Pie Scientific). For all grids described in this manuscript, an FEI Vitrobot IV (ThermoFisher) was equilibrated to 4 °C at 100% relative humidity. Grids were blotted at blot force 3 for 4 seconds and plunged into liquid ethane cooled by liquid nitrogen for vitrification.

Movies were collected on a Titan Krios G2 transmission electron microscope (ThermoFisher) at 300 keV fitted with an FEI Falcon 4 direct electron detector (ThermoFisher). For the BRISC(FL) and BRISC-FX-171-C, data were collected with a 10 eV Selectris energy filter (ThermoFisher).

For the BRISC(FL) dataset, 14,573 movies were collected using EPU automated acquisition software (v3.5.1) in counting mode. A dose per physical pixel per second of 7.97 resulting in a dose of 40.46 e^-^/Å^2^, fractionated across 855 hardware frames. These were grouped into 40 frames, resulting in a dose per frame of 1 e^-^/Å^2^, and a final pixel size of 0.74 Å/pixel. Five exposures were taken per hole, at a magnification of 165,000x, with the defocus values ranging from −0.9 μm to −2.7 μm.

For the BRISC-FX-171-C dataset, 16,750 movies were collected using EPU automated acquisition software in counting mode. A dose per physical pixel per second of 5.14 resulting in a dose of 34.97 e^-^/Å^2^, fractionated across 826 hardware frames. These were grouped into 44 frames, resulting in a dose per frame of 0.8 e^-^/Å^2^, and a final pixel size of 0.71 Å/pixel. Three exposures were taken per hole, at a magnification of 165,000x, with the defocus values ranging from −1.6 μm to −2.5 μm.

For the BRISC-JMS-175-2 dataset, 7,771 movies were collected using EPU automated acquisition software in counting mode. A dose per physical pixel per second of 4.92 resulting in a dose of 40.01 e^-^/Å^2^, fractionated across 1,428 hardware frames. These were grouped into 40 frames, resulting in a dose per frame of 0.99 e^-^/Å^2^, and a final pixel size of 0.82 Å/pixel. Three exposures were taken per hole, at a 96,000x magnification, with the defocus values ranging from −1.7 μm to −3.1 μm. More detailed data acquisition parameters are in **Extended Data Table 1**.

### Image processing

Image processing was carried out using RELION (v.3.0 and v.3.1)^64,65^. Motion correction was performed using RELION’s own implementation of the MotionCor2 algorithm^66^. Contrast transfer function (CTF) was estimated using CTFFIND (v4.1.14)^67^ for the BRISC(FL) and BRISC-FX-171-C datasets and using gCTF (v1.18)^68^ for the BRISC-JMS-175-2 dataset. Motion correction and CTF estimation were performed on-the-fly^69^.

**Extended Data Figs. 2e-g** outlines the data processing pipeline for the BRISC(FL) dataset. Particles were picked using crYOLO (v1.6.1)^70^ using a model trained on 16 micrographs. Particles coordinates were imported into RELION (v3.1.1) and 1,933,988 particles were extracted with a box size of 480 pixels. The particle stack was imported into cryoSPARC (v4.2.1)^71,72^ and subjected to two rounds of reference-free 2D classification. 148,596 particles were selected from high quality 2D class averages for *ab initio* reconstruction. Four of the six *ab initio* models corresponded to a BRISC “monomer” and were selected for heterogeneous refinement. The best class (containing 34% of the particles) was further refined using non-uniform refinement and global CTF refinement with beam tilt and beam trefoil fitted. To generate a map for the BRISC dimer complex, the class with additional density from *ab initio* reconstruction (32,283 particles) was refined using homogeneous refinement followed by non-uniform refinement and global CTF refinement with beam tilt and beam trefoil fitted. The same particles were also refined with C2 symmetry applied during non-uniform refinement. The final resolutions of both maps were determined using the gold-standard Fourier shell correlation (FSC) criterion (FSC=0.143) with FSC curves generated using the PDBe FSC server (EMDB). Local resolutions were determined using the local resolution implementation in cryoSPARC and visualised in ChimeraX (v1.2.3)^73^. To visualise the Euler angular distribution, the csparc2star.py and star2bild.py pyem scripts were used^74^. To model BRISC in the monomer and dimer conformations, one or two BRISC models with SHMT2 removed (PDB: 6H3C) were rigid body fitted using Chimera (v1.12)^75^ and visualised using ChimeraX (v1.2.3).

Data processing for the BRISC*Δ*N*Δ*C-FX-171-C dataset is outlined in **Extended Data Fig. 4b**. Briefly, a model was trained using crYOLO (v.1.6.1)^70^ using particles picked from 10 micrographs, and this model was used to pick 2,458,785 particles which were imported and extracted using RELION (v3.1.1). Particles were extracted with a box size of 192 pixels and a binning factor of two. Particles were subjected to one round of reference-free 2D classification. A BRISC-JMS-175-2 map was low-pass filtered and used as an initial model for 3D classification with no symmetry applied. The two best classes (632,988 particles) were selected for 3D refinement, post-processing and three rounds of Bayesian particle polishing and CTF refinement, resulting in a final map at 3.02 Å. To improve the density around the small molecule binding site, a mask was applied during refinement to one half of the map (**Extended Data Fig. 4g**). The resolution of the map improved to 2.8 Å overall, and 2.7 Å around the BLUE binding site.

A schematic (**Extended Data Fig. 4e**) details the data processing pipeline for the BRISC*Δ*N*Δ*C-JMS-175-2 dataset. In summary, particle picking was performed using crYOLO (v.1.6.1)^70^. A model was trained from manually picking 14 micrographs. The trained model picked 1,616,457 particles, for which the coordinates were imported into RELION (v3.1.1) for extraction with a box size of 176 pixels and a binning factor of two. Two rounds of reference-free 2D classification were used to remove junk particles. A reference model from a previous BRISC-JMS-175-2 dataset was applied during 3D classification of 1,011,924 particles with no symmetry applied. 371,872 particles were selected from three classes and re-extracted with a box size of 352 pixels. After 3D refinement and post-processing, a reconstruction of a BRISC dimer complex was achieved at 3.98 Å. Iterative rounds of per-particle CTF refinement and Bayesian polishing resulted in an improved final map at 3.32 Å. To further improve the density around the small-molecule binding site, a mask was applied during 3D refinement, encompassing only the better resolved half of the map (**Extended Data Fig. 4i**). This improved the density for “half” of the structure, resulting in a 3.2 Å map. Final resolutions were determined using the gold standard FSC criterion (FSC = 0.143). Local resolution estimation was carried out using the RELION local resolution feature.

### Model building and refinement

Atomic models of the BRISC dimer in complex with either JMS-175-2 or FX-171-C were built using high resolution cryo-EM maps. A preliminary model of the human BRCC36-Abraxas2 super dimer was acquired from our previous BRISC-SHMT2 model (PDB: 6R8F)^20^ with BRCC45 and SHMT2 removed. The BRCC36-Abraxas2 super dimer was rigid-body fitted into the cryo-EM density using UCSF Chimera^75^ and manually modelled into the BRISC-FX-171-C map using Coot^76,77^. The super dimer was duplicated, rigid-body fitted, and manually modelled into the BRCC36-Abraxas2 in the opposite side of the map. A model for human BRCC45 and MERIT40 was acquired from a previous BRISC-SHMT2 model (PDB: 6H3C)^21^ and rigid-body fitted into the cryo-EM density. The BRCC45 N-termini (residues 1-275) were manually modelled using Coot, but due to the lower resolution of the map beyond the UEV-M domain and for MERIT40, these regions were rigid-body fitted into the density based on previous BRISC-SHMT2 structures^20,21^. The BRCC45-MERIT40 arms were duplicated, rigid-body fitted, and modelled into the density corresponding to the second BRISC molecule. The side chain atoms for BRCC45 (residues 275-383) and MERIT40 were set to zero occupancy due to lower resolution of the EM maps in these regions. Small molecule chemical structures were generated in ChemDraw (PerkinElmer), and PDB and CIF files were created using the PRODRG2 server^78^ or eLBOW^79^. FX-171-C compounds were manually fit into the density using UCSF Chimera and refined using COOT Real Space Refine. The model was refined against the BRISC-FX-171-C map using Phenix real-space refinement (v1.20)^80^. To build the BRISC-JMS-175-2 structure, FX-171-C was removed from the model and replaced with JMS-175-2. The model was rigid-body fitted into the density for a BRISC-JMS-175-2 cryo-EM map and subjected to iterative rounds of manual building in Coot. The BRISC-JMS-175-2 model was refined using Phenix real-space refinement (v1.20).

### Sequence alignments, structure visualisation, and analysis

Multiple sequence alignments were performed using MUSCLE^81^ and edited using ALINE (v1.0.025)^82^. Electron microscopy maps and structure models were visualised in UCSF Chimera (v1.12.0)^75^ and ChimeraX (v1.2.3)^73^.

### Native mass spectrometry

BRISC(FL) at 10 μM was mixed with 1 mM inhibitor (JMS-175-2 or FX-171-C) or DMSO (2.5%) and incubated on ice for 30 minutes. Samples were buffer exchanged into 500 mM ammonium acetate using Zeba Spin 7K MWCO desalting columns (ThermoFisher Scientific). Samples were analysed by nanoelectrospray ionisation MS using a quadrupole-orbitrap MS (Q-Exactive UHMR, ThermoFisher Scientific) using gold/palladium coated nanospray tips prepared in-house. The MS was operated in positive ion mode using a capillary voltage of 1.5 kV, capillary temperature of 250 °C and S-lens RF of 200 V. In-source trapping was used with a desolvation voltage of −200 V for 4 μs. Extended trapping was not used. The quadrupole mass range was 2000-15000 m/z. Nitrogen gas was used in the HCD cell with a trap gas pressure setting of 5. Orbitrap resolution was 6250, detector *m/z* optimisation was low. Five microscans were averaged and an AGC target of 2 x10^5^ was used. Mass calibration was performed by a separate injection of sodium iodide at a concentration of 2 μg/μL. Data processing was performed using QualBrowser (v4.2.28.14) and deconvoluted using UniDec^83^.

### Hydrogen-deuterium exchange mass spectrometry

HDX-MS experiments were carried out using an automated HDX robot (LEAP Technologies, USA) coupled to an M-Class Acquity LC and HDX Manager (Waters Ltd., UK). For differential HDX-MS of BRISC in the absence and presence of inhibitor, 5 μM BRISC*Δ*N*Δ*C was mixed with 500 μM FX-171-C or DMSO (2.5% v/v) in buffer containing 25 mM HEPES pH 7.5, 150 mM NaCl, 1 mM TCEP. The BRISC-inhibitor or BRISC-DMSO mix was incubated on ice for 30 minutes prior to deuterium exchange reactions. For labelling, 5 μL BRISC-inhibitor/DMSO mix was diluted in 95 μL deuterated buffer (50 mM potassium phosphate, 200 mM NaCl, pH 7.5) and incubated at 4°C for 0 seconds, 0.5, 1, 10 or 60 minutes. The sample was quenched by adding quench buffer (50 mM potassium phosphate, pH 2.1) at a 1:1 ratio and dropping the temperature to 0 °C. 50 μL of quenched sample was passed through an immobilised pepsin column (AffiPro, Czech Republic) at 115 μL/min and trapped on a VanGuard Pre-column Acquity UPLC BEH C18 (1.7 µm, 2.1 mm × 5 mm, Waters Ltd., UK) for 3 min in 0.3% v/v formic acid in water. The resulting peptic peptides were transferred to a C18 column (75 µm × 150 mm, Waters Ltd., UK) and separated by gradient elution of 0–40% MeCN (0.1% v/v formic acid) in H2O (0.3% v/v formic acid) over 7 min at 40 µl min^−1^. Trapping and gradient elution of peptides was performed at 0 °C. The HDX system was interfaced to a Synapt G2Si mass spectrometer (Waters Ltd., UK). HDMSE and dynamic range extension modes (Data Independent Analysis (DIA) coupled with IMS separation) were used to separate peptides prior to CID fragmentation in the transfer cell. HDX data were analysed using PLGS (v3.0.2) and DynamX (v3.0.0) software supplied with the mass spectrometer. Restrictions for identified peptides in DynamX were as follows: minimum intensity: 10000, minimum products per MS/MS spectrum: 3, minimum products per amino acid: 0.3, maximum sequence length: 18, maximum ppm error: 10, file threshold: 8/9. Following manual curation of the data, Woods and individual uptake plots were generated using Deuteros 2.0^84^. A summary of the HDX-MS data, as recommended by reported guidelines^85^, is shown in **Extended Data Table 2**.

### Dianthus Spectral Shift binding assay

Site-specific labelling of His-tagged BRISC was performed using a RED-tris-NTA 2nd generation labelling kit (NanoTemper Technologies) in buffer containing 25 mM HEPES pH 7.5, 150 mM NaCl, 1 mM DTT, and 0.005% Tween-20. 100 nM His-BRISC was incubated with 25 nM RED-tris-NTA dye and prepared according to the recommended protocol from NanoTemper. To measure the affinity of the interaction between BRISC and SHMT2, 12.5 nM labelled BRISC was mixed with SHMT2(A285T) in a 16-point, 2-fold dilution series from 46 μM to 1.4 nM. To measure the effect of the compounds on the BRISC-SHMT2 interaction, NTA-labelled His-BRISC was incubated with 1% DMSO, or 100 μM compound (1% DMSO) for 15 minutes at RT. The BRISC-compound mix was then incubated with SHMT2(A285T) for 30 minutes at 25 °C. The reactions were carried ot in 384-well Dianthus microplates (Nanotemper Technologies) with a 20 μL reaction volume. The measurements were performed using auto-excitation on a Dianthus NT.23 instrument at 25 °C using DI.Control software (v2.1.1) (Nanotemper Technologies). Data were analysed using DI.Screening Analysis software (v2.1.1) (Nanotemper Technologies) and plotted in GraphPad Prism (v10.1.0). *K*_D_ values were determined using a GraphPad Prism built-in equation for total binding (one-site).

### Cell lysis, immunoprecipitation, and immunoblotting

Immortalised breast epithelial cell line MCF10A were cultured in 10 cm plates with Dulbecco’s Modified Eagle Medium (DMEM) + F-12 (1:1) supplemented with 5% horse serum, 100 ng/mL cholera toxin, 10 μg/mL insulin, 20 ng/mL epidermal growth factor (EGF) and 0.5 μg/mL hydrocortisone until >80% confluence. 1.2 x10^5^ cells were then seeded into 6 cm plates. Cells were left to reach ultra-confluence. Once confluent, cells were washed with phosphate buffered saline (PBS) and starved for 30 minutes in 2 mL DMEM + F-12 (1:1) supplemented with 1% horse serum. After 30 minutes cells were pre-treated with either DMSO (0.1%) (control and IFN only) or BRISC inhibitor (2.5 μM) for 15 minutes. After 15 minutes, hIFN2α (75 ng/mL) was added to the media and mixed it well. Cells were treated for indicated time or otherwise 1 hour (without inhibitors) or 4 hours (with BRISC inhibitors). Cells were washed with ice cold PBS and scraped with cell scrapers into PBS and centrifuged at 1,200 rpm for 10 minutes. Cell pellets were either resuspended into radioimmunoprecipitation assay (RIPA) buffer supplemented with complete ethylenediaminetetraacetic acid (EDTA) free protease inhibitor cocktail and 25 U/mL benzonase for immunoblotting or in immunoprecipitation buffer (100 mM NaCl, 0.2% Igepal CA-630, 1 mM MgCl_2_, 10% glycerol, 5 mM NaF, 50 mM Tris-HCl, pH 7.5), supplemented with complete EDTA free protease inhibitor cocktail and 25 U/mL benzonase for co-immunoprecipitation. Lysates were mixed with 2X sample buffer for gel loading prior to Western blot analysis. Co-immunoprecipitation was performed using Anti-Flag M2 Affinity Gel beads and eluted with 0.2 M Glycine.

### Real-time quantitative PCR

1×10^5^ MCF10A cells were cultured as previously described and seeded into 6 well plates. Cells were left to reach ultra-confluence. Once confluent, cells were washed with PBS and starved for 30 minutes in 2 mL DMEM + F-12 (1:1) supplemented with 1% horse serum. After 30 minutes cells were pre-treated with either DMSO (0.1%) (Control and IFN only) or BRISC inhibitor (2.5 μM) for 15 minutes. After 15 minutes, hIFN2α (75 ng/mL) was added to the media and mixed. Cells were treated for 4 hours. Cells were washed with ice cold PBS and scraped with cell scrapers into 350 μL RNAlater buffer and RNA was isolated using RNeasy mini kit and treated with on column DNase. Isolated RNA was quantified using a Nanodrop spectrophotometer. RNA was converted into cDNA and the complementary DNA was used for real-time quantitative PCR analysis of expression of interferon stimulated genes (ISG15, IFIT1, IFIT2, IFITM1, CXCL10 and 18s rRNA) using an Applied Biosystems Quantstudio 6 RT-PCR system. All experiments were carried out in triplicate.

### IFNAR1 flow cytometry

To study the direct effect of BRISC DUB inhibitors, IFNAR1 surface levels were measured using flow cytometry (FACS) analysis. For this purpose, MCF10A WT and MCF10A BRCC45 KO cells expressing BRCC45 WT or BRCC45 R137A mutant were used. Cells were pre-incubated with 5 μM FX-171-C for 30 minutes prior to the addition of 50 ng/mL of hIFN-Iα for various time points (45 minutes and 90 minutes). After IFN treatment, cells were dissociated from plate using cell dissociation Buffer (ThermoFisher Scientific, #13150016) and were stained with LIVE/DEAD™ Fixable Violet Dead Cell Stain Kit, for 405 nm excitation (ThermoFisher Scientific, #L34963) for 30 minutes, as per the kit protocol. The cells were then washed using FACS buffer (DPBS, 1% BSA, 2 mM EDTA, 0.09% sodium azide) and transferred to 96-well round bottom plates. Resuspended cells were divided into three repeats for each sample, each was surface stained for 1 hour at 4 °C with: 1. Human IFN-alpha/beta R1 Antibody (R&D systems, #MAB245), 2. Mouse IgG1 Isotype Control (R&D systems, #MAB002), 3. FACS buffer (control cells). Following incubation, cells were washed once with FACS buffer and were labelled for 30 minutes at 4 °C with either Biotin-SP-conjugated AffiniPure Donkey Anti-Mouse IgG (H+L) (Jackson Immunoresearch Laboratories, Inc.) for samples 1 and 2 or with FACS buffer for sample 3. Then, cells were washed twice with FACS buffer, and were labelled for 15 minutes at 4 °C with either R-Phycoerythrin-conjugated Streptavidin (Jackson Immunoresearch Laboratories, Inc.) for samples 1 and 2 or with FACS buffer for sample 3. Cells were washed 3 times with FACS buffer and were analysed by FACS (MACSQuant, Miltenyi Biotec). For each sample from group 1 the Geo mean fluorescence intensity (MFI) of its isotype sample was reduced. IFNAR1 cell surface percentage was calculated by 100*(MFI treated sample/MFI of untreated sample). Statistical analysis was performed by one-way and two-way ANOVA.

### THP-1 *in vitro* experiments

THP-1-Dual^TM^ cells (InvivoGen), derived from a human monocytic cell line, isolated from a male patient with acute monocytic leukaemia, were cultured as per manufacturer’s guidance. Cells were seeded at 1×10^6^ cells/mL in RPMI1640 media containing 10% fetal bovine serum (FBS), 1% penicillin-streptomycin (PS) (Gibco Laboratories) with DMSO (0.1%) (control) or in media containing JMS-175-2, FX-171-C, FX-171-A, or AP-5-145 at 4 µM for 16 hours with and without IFNα2 (25 ng/mL), LPS (100 ng/mL) (Merck Life Science), ODN 2216 (1 µM) (Miltenyi Biotec), or Poly:IC (1 µg/mL) (InvivoGen). Tofacitinib (Cambridge Biosciences) at 0.4 µM was used for comparison against a known JAK inhibitor. Cells were centrifuged at 300 g for 5 minutes. Supernatant was removed and subjected to simultaneous study of the NF-κB pathway, by monitoring the activity of secreted alkaline phosphatase reporter (SEAP), and the IRF pathway, by assessing the activity of a secreted Lucia luciferase using QUANTI-Blue™ Solution and QUANTI-Luc™ 4 Lucia/Gaussia (InvivoGen), respectively, as per manufacturer’s instructions. Pelleted cells were used for FACS analysis using IFNα/β R1 APC-conjugated antibody or isotype control (Biotechne, FAB245A and IC002A), or unstained staining procedure with and without Zombie Green^TM^ (Biolegend). Samples were analysed using CytoFlex LX (Beckman) (50,000 live cells per condition). Median fluorescence intensity for APC channel gated on single cell live cells was analysed and compared as a percentage to control (no stimulation condition).

### Tandem ubiquitin binding entity (TUBE) pull-down assay

The TUBE pull-down experiment was performed according to the manufacturer’s (UM401: TUBE 1 Agarose; LifeSensors) protocol with modifications. MCF10A cells (∼20 x10^6^) were pre-treated with the BLUE inhibitor panel (5 µM) or DMSO vehicle control for 1 hour followed by addition of recombinant IFNα2 (50 ng/mL, 592702, BioLegend). Following 1 hour incubation with IFNα2, cells were lysed in cell lysis buffer (50 mM Tris-HCl, pH 7.5, 0.15 M NaCl, 1 mM EDTA, 1% NP-40, 10% glycerol, 10 mM NEM). Cell lysates were clarified by centrifugation at 15,000 g for 10 minutes at 4 °C. Clarified lysates were pre-cleared with control uncoupled agarose beads (UM400, LifeSensors) for 30 minutes on a rotator at 4 °C followed by centrifugation at 5,000 g for 2 minutes. Equal amount of pre-cleared lysates from individual samples were incubated with 20 µL Agarose-TUBE beads (pre-equilibrated with 1X Tris-buffered saline (TBS)) overnight on a rotator at 4 °C. Beads were collected by centrifugation at 5,000 g for 2 minutes at RT followed by four stringent washes with 1X TBST (TBS, 0.1% Tween-20). TUBE-pulldown proteins were eluted by boiling Agarose-TUBE beads in 2X LDS sample buffer supplemented with 200 µM DTT for 15 minutes. Eluted proteins were resolved on NuPAGE™ 4–12% Bis–Tris gel followed by Western blotting using anti-IFNAR1 antibody (ab124764, Abcam).

### PBMCs *in vitro* experiments

Peripheral blood mononuclear cells (PBMCs) from healthy and SSc donors were separated using density gradient method (Leucosep™, Greiner Bio-One International) from EDTA anticoagulated peripheral blood; isolated cells were washed twice by PBS. PBMCs were then incubated in RPMI1640 media containing 10% FBS, 1% PS with DMSO (0.1%) (CTR) or in media containing JMS-175-2, FX-171-C, or AP-5-145 at 2 µM for 16 hours. For healthy PBMCs, cells were stimulated with 20 ng/mL of IFN (+/-compounds or DMSO). For SSc PBMCs, cells were in basal conditions with no additional IFN stimulation. Cells were centrifuged at 300 g for 10 minutes. Supernatant was removed and subjected to R&D Systems DY266 Human CXCL10/IP-10 DuoSet ELISA (Biotechne) in duplicates, according to manufacturer’s guidance. Concentration of CXCL10 was determined according to a 4-parameter logistic regression standard curve. RNA was extracted from cell pellets using TRIzol™ (ThermoFisher Scientific, Waltham, MA) and processed using Quick-RNA Miniprep Kit (Zymo Research) as per the manufacturer’s instruction.

### Gene expression analysis

For Type I IFN signalling analysis, the human IFN I RT2 Profiler PCR Array (Qiagen, PAMM-016ZE) was performed on available RNA from healthy (n=3) and SSc (n=9) donors. RNA was converted to cDNA using RT2 First Strand Kit (Qiagen). Next, the cDNA was mixed with RT2 SYBR Green Mastermix (Qiagen, Venlo). Successful PCR performance was confirmed by assay tests for genomic DNA contamination, RNA sample quality, as well as single melt curve determination and <35 Ct for each gene, leading to validated gene expression of 67 genes. Relative expression for each gene was determined, firstly by ΔCT calculated using the geometric mean of 4 housekeeping genes (ACTB, GAPDH, HPRT1, RPLP0), followed by 2^(-ΔCT), and expressed as fold change to control, according to manufacturer’s guidance. IFN2*α*-induced ISGs in healthy donors were determined using a Student’s t-test (two-tail distribution and equal variances) on the triplicate fold change in 2^(-ΔCT) values between treatment and control group (AP-5-145+IFN2*α* v DMSO only conditions; 34 genes p<0.05 for n=3 donors). Heatmaps were generated using the “pheatmap” function in R (version 4.0.2) and RStudio (version 1.3.1093) applying row scaling to standardise the expression data across samples (R core team 2020. R Foundation for Statistical Computing). The IFN2α inducible genes determined in healthy donors (n=34) were included, and donors were combined into treatment groups AP-5-145, JMS-175-2 and FX-171-C. To account for Type I IFN signalling heterogeneity across samples, the mean 2^(-ΔCT) for each gene for AP-5-145 treated condition from all donors (healthy, n=3 or SSc, n=9) was used to determine relative gene expression, illustrated as a fold change and visualised in log2. The Type I IFN signalling array data for the patient samples is is included in the **Supplementary Information**.

For composite ISG score gene expression analysis, RNA was reverse transcribed using the high-capacity cDNA synthesis kit (Applied Biosystems). qRT-PCR were performed using SyBr Green PCR kit (ThermoFisher Scientific) with primers specific for CXCL10 (Forward; TCCAGTCTCAGCACCATGAA Reverse; AGGTACTCCTTGAATGCCACT), MX1 (Forward; CGACACGAGTTCCACAAATG Reverse; AAGCCTGGCAGCTCTCTACC), IFIT1 (Forward; GACTGGCAGAAGCCCAGACT Reverse; GCGGAAGGGATTTGAAAGCT), ISG15 (Forward; GTGGACAAATGCGACGAACC Reverse; ATTTCCGGCCCTTGATCCTG), and GAPDH (Forward; ACCCACTCCTCCACCTTTGA Reverse; CTGTTGCTGTAGCCAAATTCGT). The data obtained was analysed according to the ΔΔ Ct method relative to GAPDH. For composite IFN score, fold change in gene expression in AP-5-145, JMS-175-2 and FX-171-C treated samples (of each gene CXCL10, MX1, IFIT1, ISG15) was calculated relative to each donor DMSO control (CTR). The composite score represents grouped analysis of combined fold changes for all 4 genes.

### Inclusion and ethics

All participants enrolled provided written informed consent according to a protocol approved by Medicine and Health Regulatory agency (STRIKE NRES-011NE to FDG, IRAS 15/NE/0211).

### Statistical analysis

Comparisons between two conditions were conducted using paired and unpaired student t-tests. Comparisons between multiple conditions were conducted using either a one-way ANOVA with Dunnett’s multiple comparisons test or two-way ANOVAs. Statistical significance was defined as a p-value less than 0.05 for all analyses. Data analysis was performed using GraphPad Prism (v9.5.1).

## Data availability

Cryo-EM maps have been deposited in the Electron Microscopy Data bank under the accession codes EMD-17980 and EMD-18009. Model coordinates have been deposited in the Protein Data Bank under the accession codes 8PVY, 8PY2. HDX data are available via ProteomeXchange (identifier: PXD044584). All unique reagents are available upon request.

## Acknowledgements

We thank R. George, T. Sun, and F. Vizeacoumar, for technical support and D. Maskell for assistance with cryo-EM data collections. Mass photometry experiments were performed at Refeyn (Oxford, UK), with support from J. Wilkinson and J. Andrecka, and in the Faculty of Biological Sciences Biomolecular Interactions Facility through Wolfson Trust-funding, in partnership with the Bragg Centre.

## Author contributions

F.C. performed mass photometry, protein production (WT and mutants), enzyme activity assays, compound and SHMT2 inhibition assays, and Spectral Shift assays. F.C. and M.W. performed negative stain electron microscopy and cryo-electron microscopy. F.C. performed the electron microscopy data processing and model building. F.C., J.R.A., and A.N.C. performed native and HDX mass spectrometry experiments. P.A.N.R. and J.M.S. performed chemical synthesis. S.B., A.D., D.N.S., R.A.G. generated inhibitor-insensitive cell lines, analysed IFN response in cells with qPCR, performed FACS and TUBE analysis. K.W., R.L.R, S.D., F.D.G., performed experiments in THP-1 cells and analysed IFN signalling in PBMCs from healthy volunteers and scleroderma patients. J.C. and L.B. performed *in vitro* IC_50_ measurements. M.A.P., A.A., and R.S.A. assembled the chemical library and performed fragmentation analysis. E.Z., A.D. and F.S. performed compound screening. L.J.C., M.F., M.W., E.Z., F.S. were involved with protein production. F.C., R.L.R., F.D.G., J.M.S., R.A.G. and E.Z. performed the majority of experimental design and data interpretation. F.C. and E.Z. wrote the first draft of the manuscript with input from R.L.R., S.B., F.D.G., J.M.S. and R.A.G..

## Extended Data Figure Legends

**Extended Data Figure 1.**
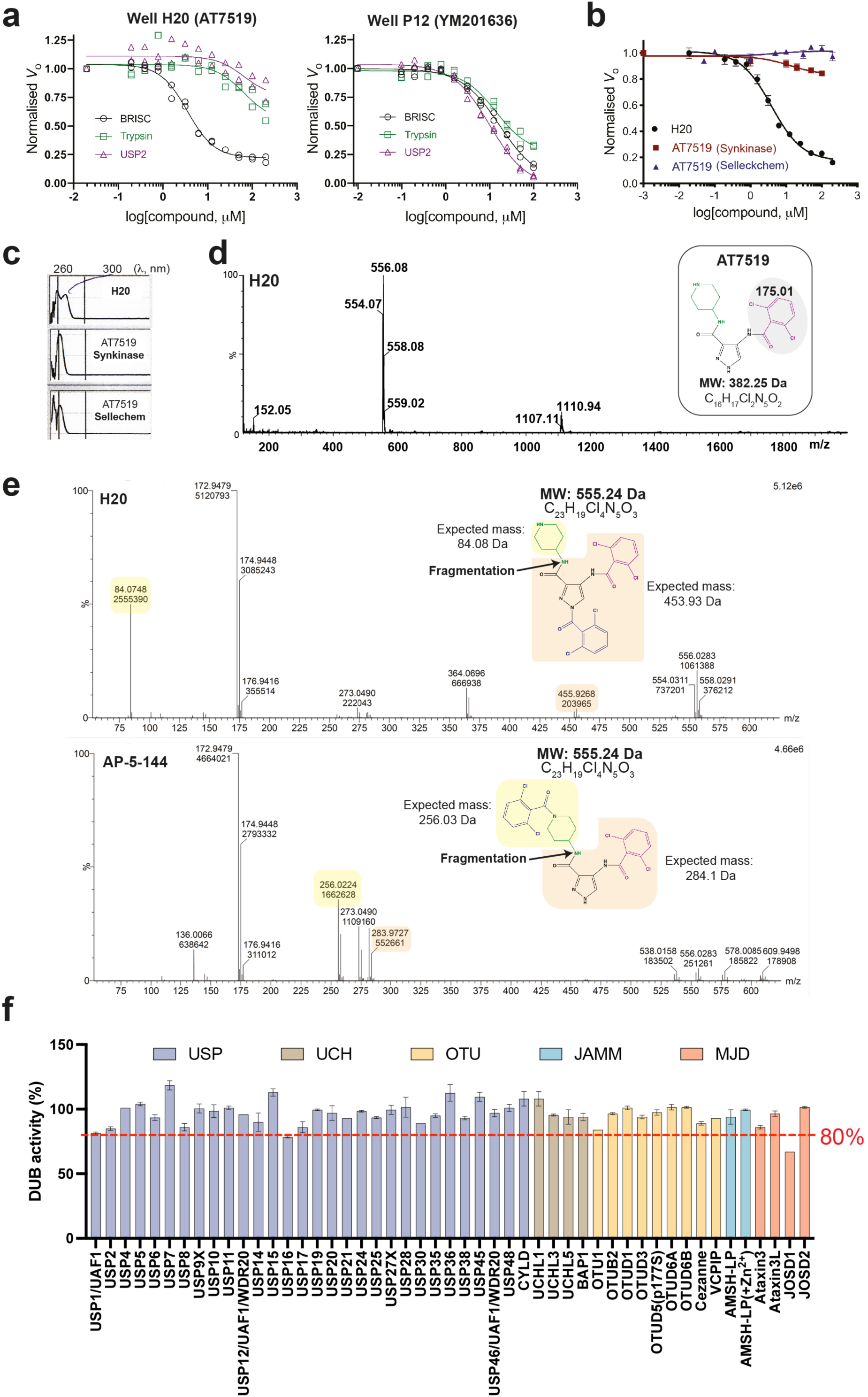
Validation of hit compounds. **a,** Dose response curves for hit compounds against BRISC (1 nM), USP2 (100 nM) and trypsin (125 nM) using the internally-quenched fluorescence di-ubiquitin assay described in Fig. 1a. Data points from two experimental replicates are plotted. **b,** Re-testing of purchased H20 hit compound presumed to be AT7519. Data points are mean ± SEM from three independent experiments. **c,** UV-vis profile of compound in well H20 and purchased AT7519 compounds from Synkinase and Selleckchem. **d,** Liquid-chromatography mass spectrometry (LC-MS) spectra of H20 compound. *Inset*, AT7519 structure. The difference between the H20 compound and AT7519 is 173 Da, which corresponds to the mass of a dichlorobenzaldehyde group. **e,** MS fragmentation analyses for H20 compound and a synthesised isomer, AP-5-144. **f,** Profiling JMS-175-2 activity (5 µM) against a panel of 48 available DUBs using a ubiquitin-rhodamine(110)-glycine enzymatic assay.

**Extended Data Figure 2.**
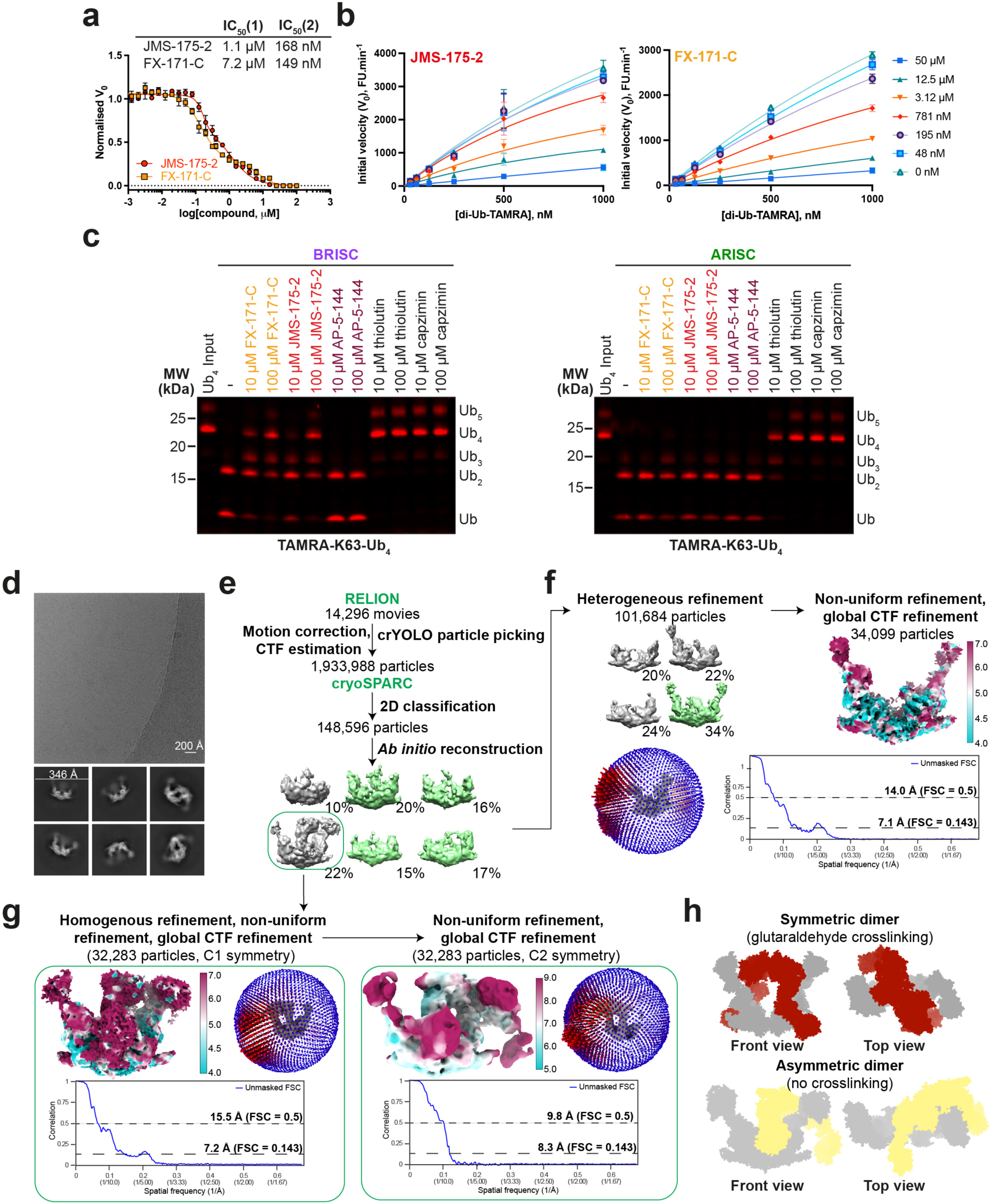
Identification of a higher-order BRISC conformation. **a,** 32-point dose-response inhibition assay with JMS-175-2 and FX-171-C, with a biphasic curve fitted. Data points are mean ± SEM of two independent experiments carried out in technical duplicate. **b,** Michaelis-Menten plots of BRISC activity against a K63-linked di-ubiquitin fluorogenic substrate with increasing concentrations of JMS-175-2 and FX-171-C. Data points are mean ± SEM of three independent experiments carried out in technical duplicate. **c,** In-gel DUB assay comparing cleavage of a TAMRA-labelled K63-linked tetraubiquitin substrate by BRISC (*left*) and ARISC (*right*) with indicated compounds. For gel source data, see **Supplementary** Figure 1. **d,** Representative micrograph (BRISC dataset) and corresponding 2D class averages generated in cryoSPARC. **e-f,** Cryo-EM processing workflow for BRISC **f,** monomer and **g,** dimer. Green indicates selected classes for 3D refinement in cryoSPARC. **f,** Final monomer cryo-EM density map coloured by local resolution and Euler angular distribution (*left*). Rod heights are proportional to the number of particles in each direction. Unmasked FSC curves with resolution calculated using the gold standard FSC cut-off at 0.143 and 0.5 frequency. **g,** Final dimer maps with C1 and C2 symmetry applied, coloured by local resolution. Euler angular distribution shown with rods, and unmasked FSC curves, as in **f**. **h,** *Top*, surface model of ARISC dimers observed in negative stain EM from grids prepared using the GraFix cross-linking method. The same conformation is reported for BRISC dimers from nsEM grids prepared using GraFix. *Bottom*, an asymmetric BRISC dimer conformation observed in cryo-EM without cross-linking.

**Extended Data Figure 3.**
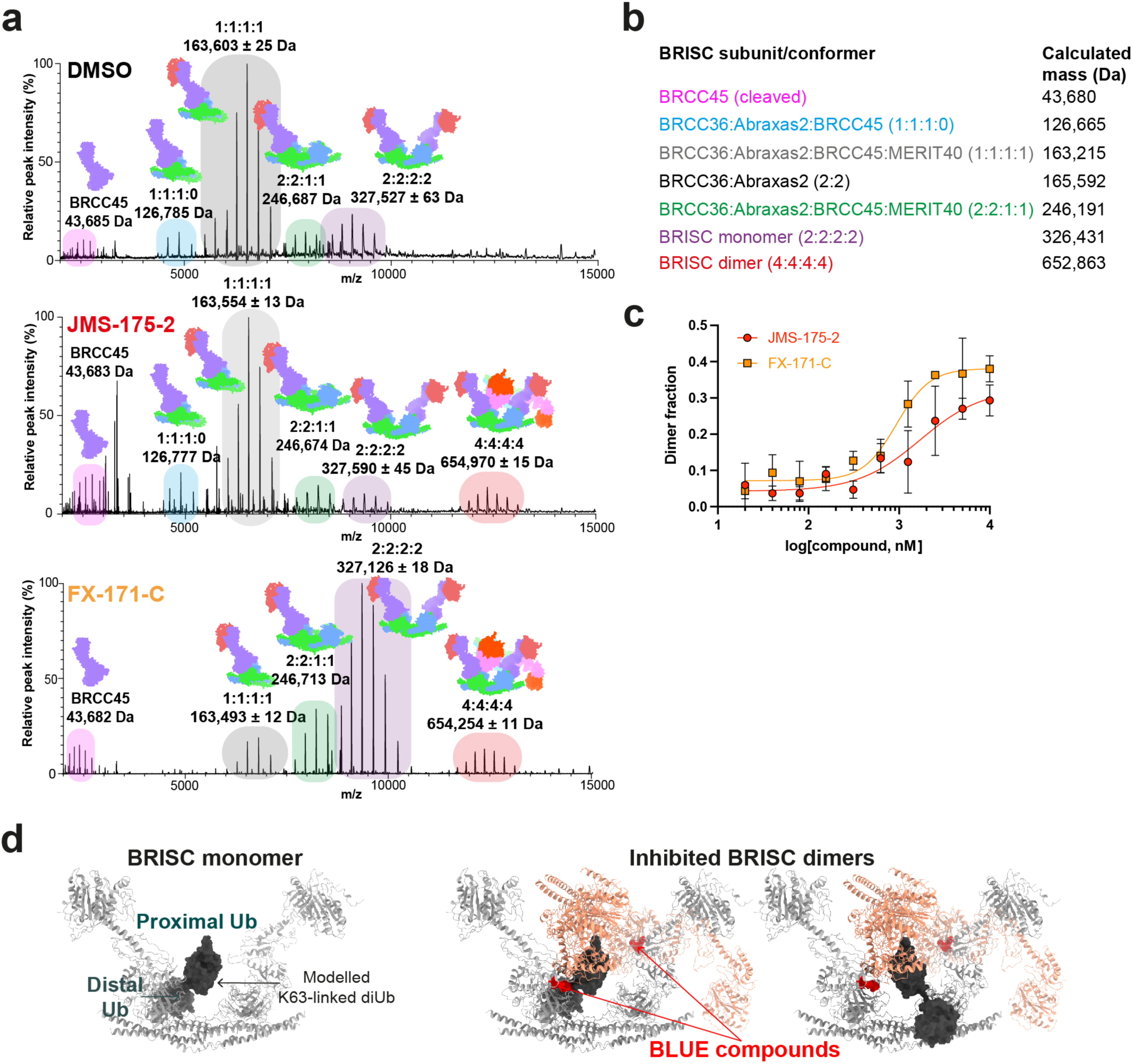
Identification of BRISC dimers in mass photometry and native mass spectrometry. **a,** Native mass spectra of BRISC mixed with DMSO (control), JMS-175-2, or FX-171-C. BRISC complexes and subcomplexes are highlighted. **b,** Table of calculated masses for different BRISC subcomplexes and super complexes. **c,** Mass photometry measurements of BRISC dimer at increasing inhibitor concentrations. Counts corresponding to BRISC dimer as a fraction of total counts are plotted. Data points are mean ± SEM from three independent experiments. **d,** *Left,* K63-linked diUb (dark grey) modelled on the MPN^+^ domain of BRCC36 in BRISC (light grey), based on the AMSH LP-diUb structure (PDB: 2ZNV). *Right,* Upon dimer formation, the second BRISC monomer sterically clashes with the proximal ubiquitin when it is bound to either BRCC36 active site.

**Extended Data Figure 4.**
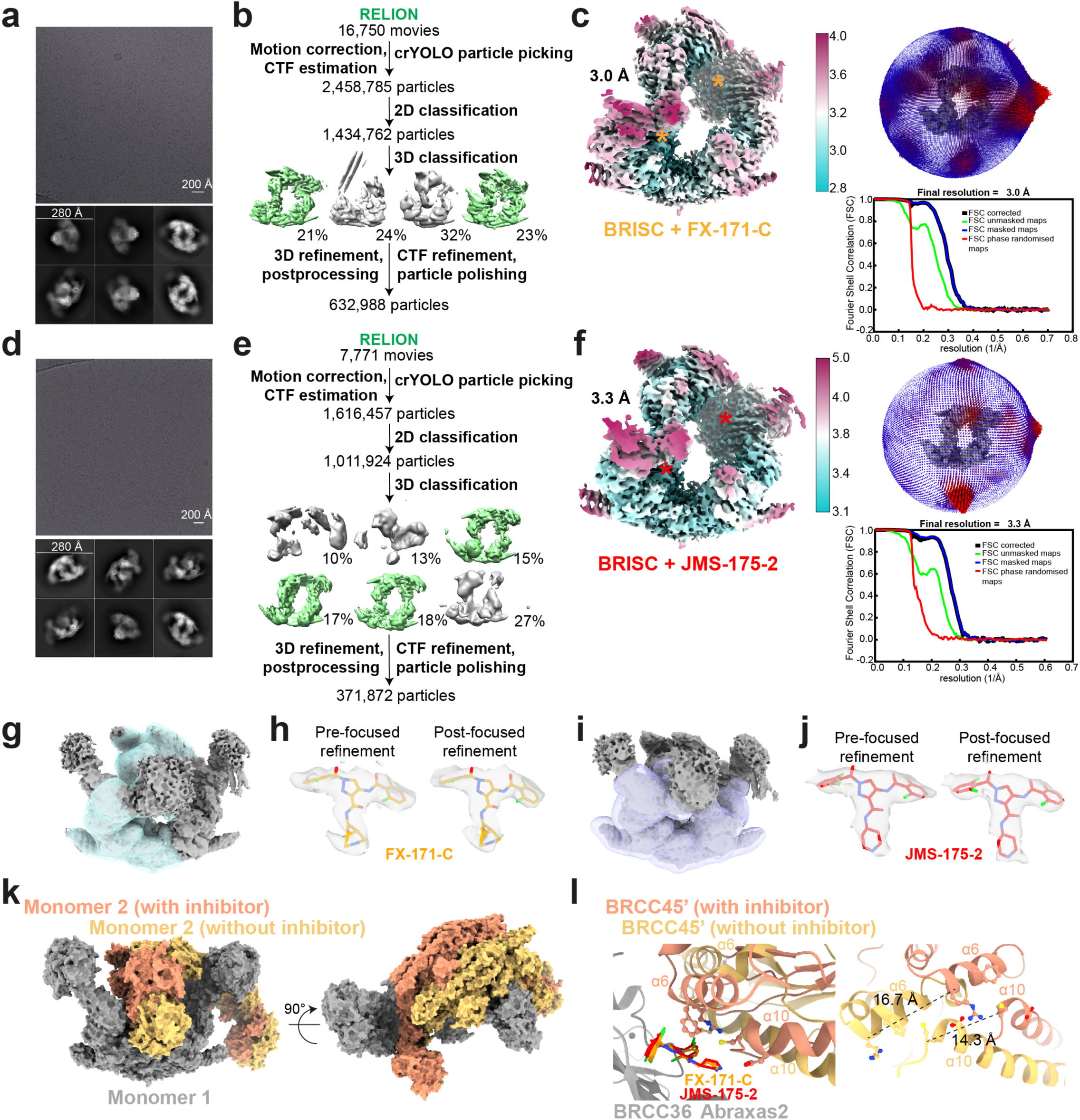
Cryo-EM processing of the BRISC-inhibitor co-complex. Figures **a-c** correspond to the BRISC-FX-171-C cryo-EM dataset. Figures **d-f** correspond to the BRISC-JMS-175-2 dataset. **a, d,** Representative micrographs and 2D class averages. **b, e,** Image processing workflow. Green maps indicate selected classes used for 3D refinement. **c, f,** *Left*, cryo-EM density maps after 3D refinement for the final reconstructions used for model building. Asterisks indicate BLUE compound binding sites. *Right*, final maps with corresponding Euler angular distribution with rod heights proportional to the number of particles in each direction. FSC curves with resolution calculated using the gold standard FSC cut-off at 0.143 frequency. **g,** Mask used for focused refinement of the BRISC-FX-171-C map. **h,** Chemical structure of FX-171-C fitted into EM density before (*left*) and after (*right*) focused refinement. Cryo-EM density visualised using the surface zone tool in ChimeraX; left, radius 2.04, right, radius 2.60. **i,** Mask applied during refinement of BRISC-JMS-175-2 map. **j,** Chemical structure of JMS-175-2 fitted into EM density before (*left*) and after (*right*) focused refinement. Cryo-EM density visualised using the surface zone tool in ChimeraX; left, radius 2.20, right, radius 2.41. **k,** Overlay of two BRISC dimers aligned on one BRISC molecule (grey) for comparison. Models are represented as surfaces. Orange, model fitted to the BRISC-FX-171-C structure shown in **c,** yellow, BRISC models rigid-body fitted in the cryo-EM density of the asymmetric dimer shown in **Extended Data Fig. 2g**. The yellow molecule is shifted relative to the orange molecule. **l,** Models described in **k,** focussed on the small molecule binding site highlighting the shift in the BRCC45’ α6 and α10 helices.

**Extended Data Figure 5.**
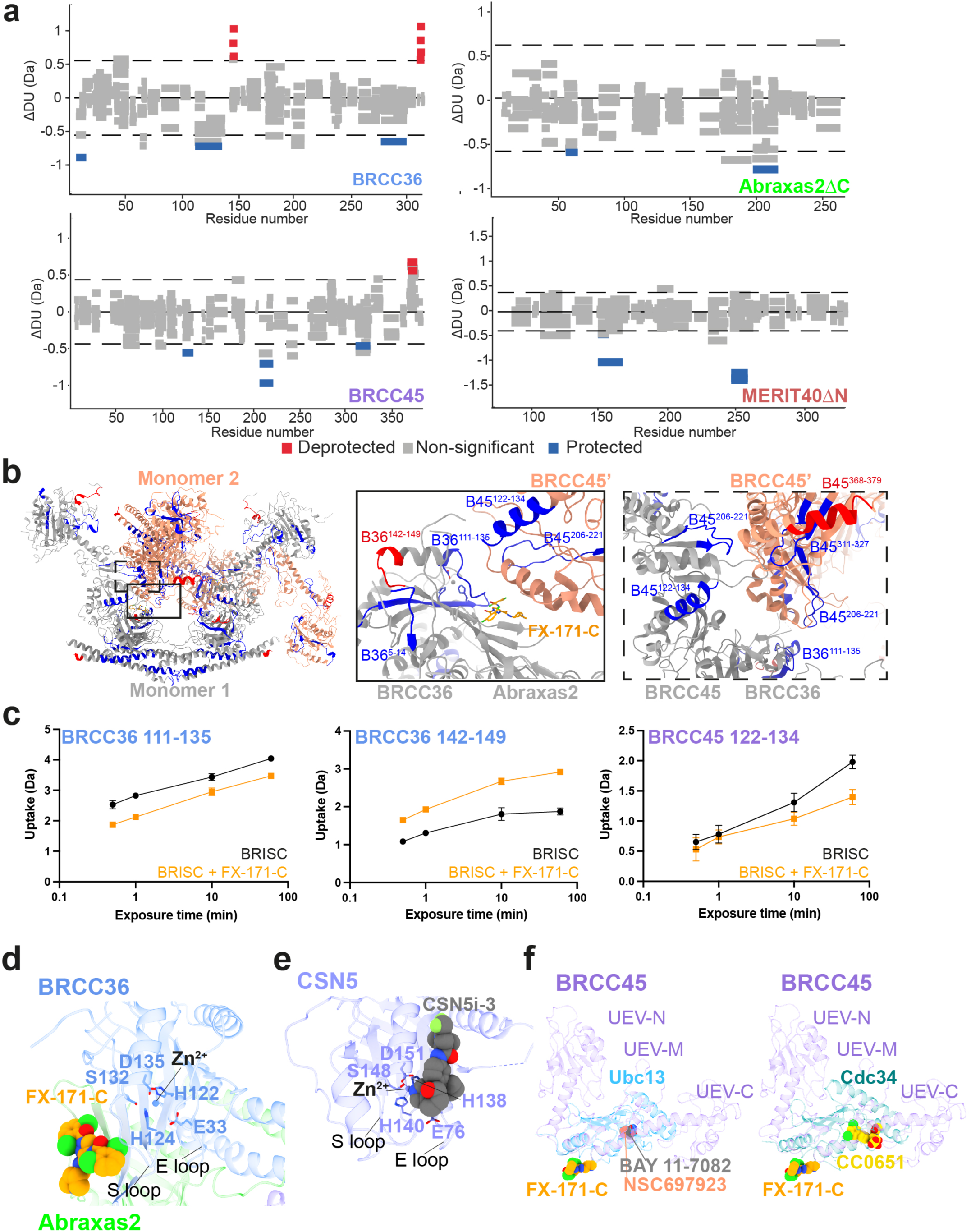
Observed changes in BRISC subunit solvent accessibility and secondary structure in the presence of FX-171-C by HDX-MS. **a,** Wood’s plots generated with Deuteros showing the differences in deuterium uptake over all four HDX timepoints from three technical replicates, comparing BRISC in the absence and presence of FX-171-C. Regions highlighted in grey indicate peptides with no significant change, calculated using a 99% confidence interval, between the two conditions. The dashed line indicates the 99% confidence limit. Peptides are coloured in red to indicate deprotection in the presence of inhibitor, and blue to indicate protection. **b,** Peptides mapped onto BRISC dimer structure, highlighting peptides near BLUE binding site and at the interface of two BRCC45 subunits. B36 = BRCC36; B45 = BRCC45. **c,** Example deuterium uptake curves in the absence and presence of FX-171-C. Data points are mean ± SEM from three technical replicates. **d,** BLUE compounds are allosteric inhibitors and do not disrupt the BRCC36 Zn^2+^ binding site. **e,** CSN5 active site in complex with inhibitor CSN5i-3 (PDB: 5J0G). f, *Left*, BRCC45 UEV-M bound to FX-171-C aligned to Ubc13 in complex with BAY 11-7082 (PDB: 4ONN) and NSC697923 (PDB: 4ONM). *Right*, BRCC45 UEV-M bound to FX-171-C aligned to Cdc34 in complex with CC0651 (PDB: 3RZ3).

**Extended Data Figure 6.**
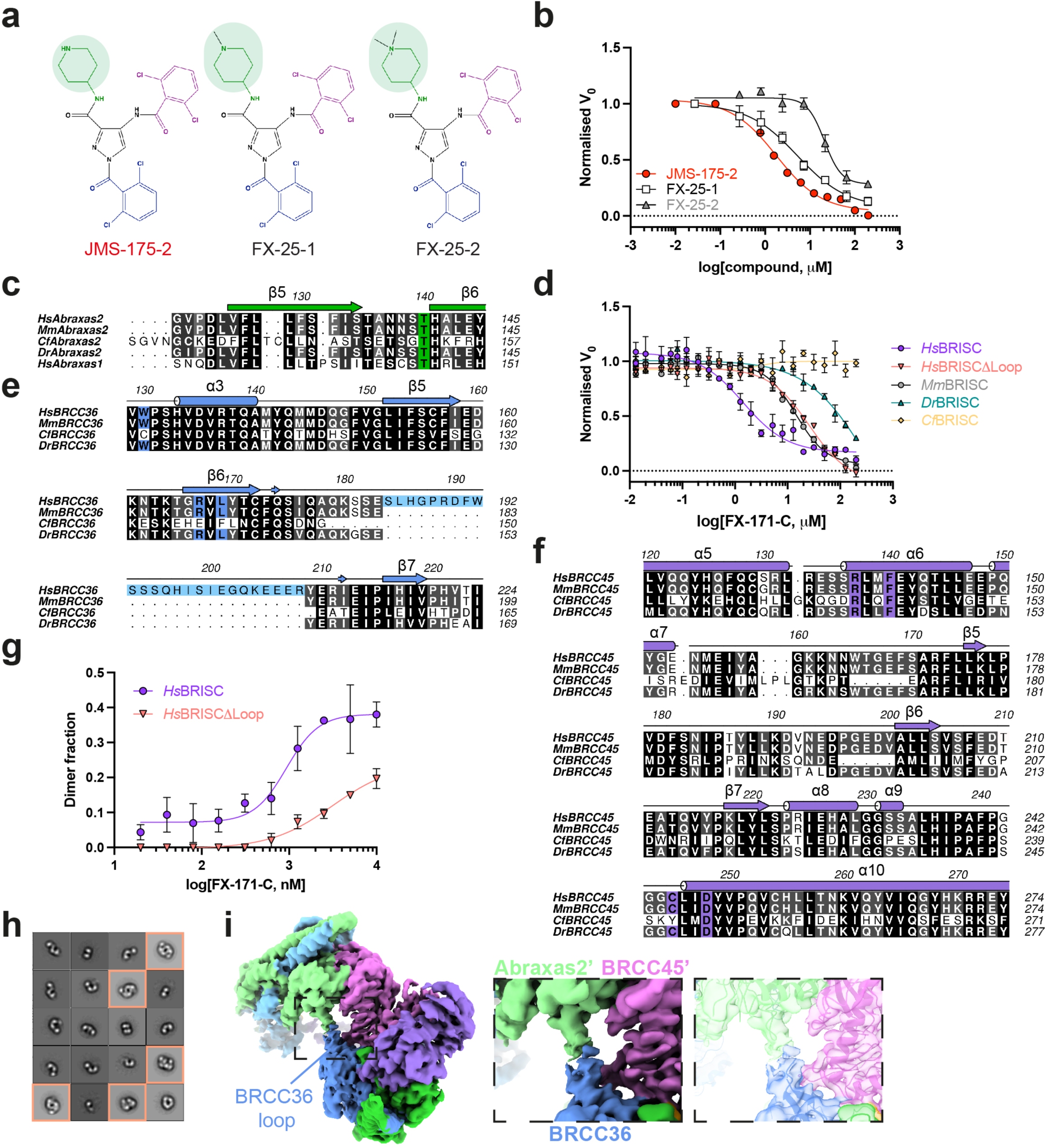
BLUE compounds are allosteric inhibitors and selective for human BRISC. **a,** Chemical structure of JMS-175-2 and analogues FX-25-1, FX-25-2, which have substitutions in the piperidine ring (highlighted in green). **b,** Dose-response inhibition of BRISC by indicated compounds. IC_50_ values: JMS-175-2 = 3.8 µM, FX-25-1 = 5.2 µM, FX-25-2 = 21 µM. Data points are mean ± SEM of three independent experiments carried out in technical duplicate. **c,** Multiple sequence alignment (black = conserved, white = not conserved) of Abraxas1 and Abraxas2 from indicated species. Coloured boxes indicate BLUE interacting residues. **d,** FX-171-C inhibition of different BRISC orthologues. *Hs - H. sapiens, Mm - M. musculus, Dr - D. rerio, Cf - C. floridanus.* Data points are mean ± SEM of three independent experiments carried out in technical duplicate. **e, f,** Multiple sequence alignment of **e,** BRCC36 and **f,** BRCC45 from indicated BRISC orthologues Residues are colored as in **c,**. **g,** Mass photometry analyses of dimer formation with FX-171-C for *Hs*BRISCΔNΔC and *Hs*BRISCΔLoop. Fraction of counts corresponding to BRISC dimer are plotted. Data points are mean ± SEM of three independent experiments. **h,** Negative stain EM 2D class averages of *Hs*BRISCΔLoop incubated with FX-171-C. 22% of particles in the 2D class averages correspond to BRISC dimers. **i,** BRISC-FX-171-C cryo-EM density map highlighting an extended loop in BRCC36 (dust cleaning size 7.1, map threshold 0.0044).

**Extended Data Figure 7.**
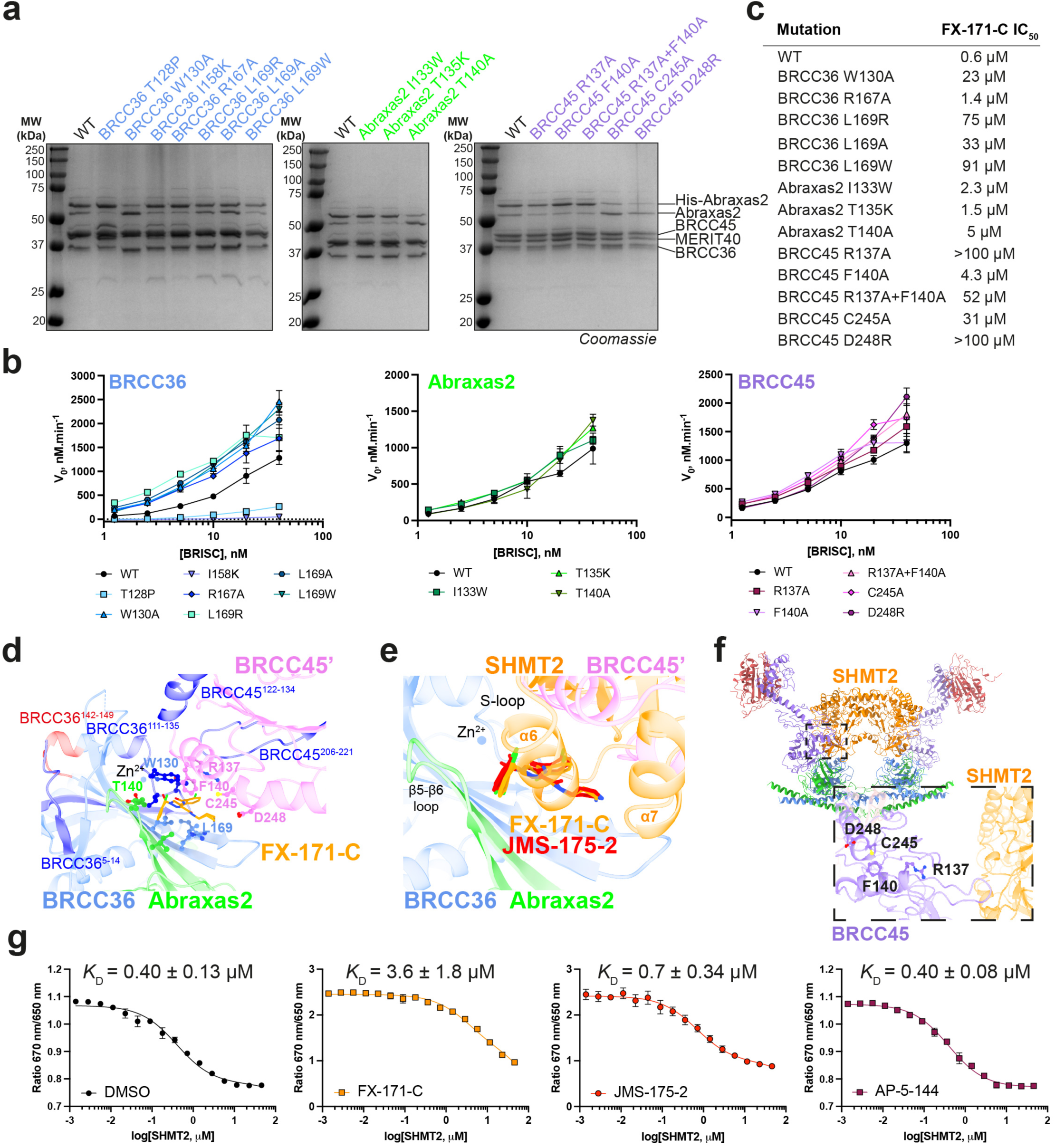
Determining the DUB activity, inhibitor sensitivity, and SHMT2 inhibition of structure-guided mutants. **a,** SDS-PAGE analysis of purified BRISC mutants. For gel source data, see **Supplementary** Figure 1. **b,** Activity of BRISC mutants against an IQF di-ubiquitin substrate. Data points are mean ± SEM of three independent experiments carried out in technical duplicate. **c,** FX-171-C IC_50_ values from inhibition assays shown in Fig. 3. **d,** Protected and deprotected peptides from HDX-MS mapped onto the FX-171-C binding site. Peptides are coloured blue to indicate protection and red to indicate deprotection, after incubation with FX-171-C. **e,** Superimposition of the SHMT2 dimer from BRISC-SHMT2 structure (PDB: 6R8F) onto BRISC-FX-171-C dimer structure. SHMT2 α6 helix clashes with the BLUE binding site. **f,** Mutated residues in BRCC45 are not in close proximity to the SHMT2 binding site in the BRISC-SHMT2 structure (PDB: 6R8F). **g,** Spectral Shift (Dianthus) assays measure the binding of SHMT2(A285T) to labelled His-BRISC in the absence and presence of compounds. *K*_D_ is calculated by plotting the ratio of the fluorescence intensities at 650 nm and 670 nm against SHMT2 concentration, with a GraphPad Prism equation for one-site total binding. Data points are mean ± SEM of three independent experiments.

**Extended Data Figure 8.**
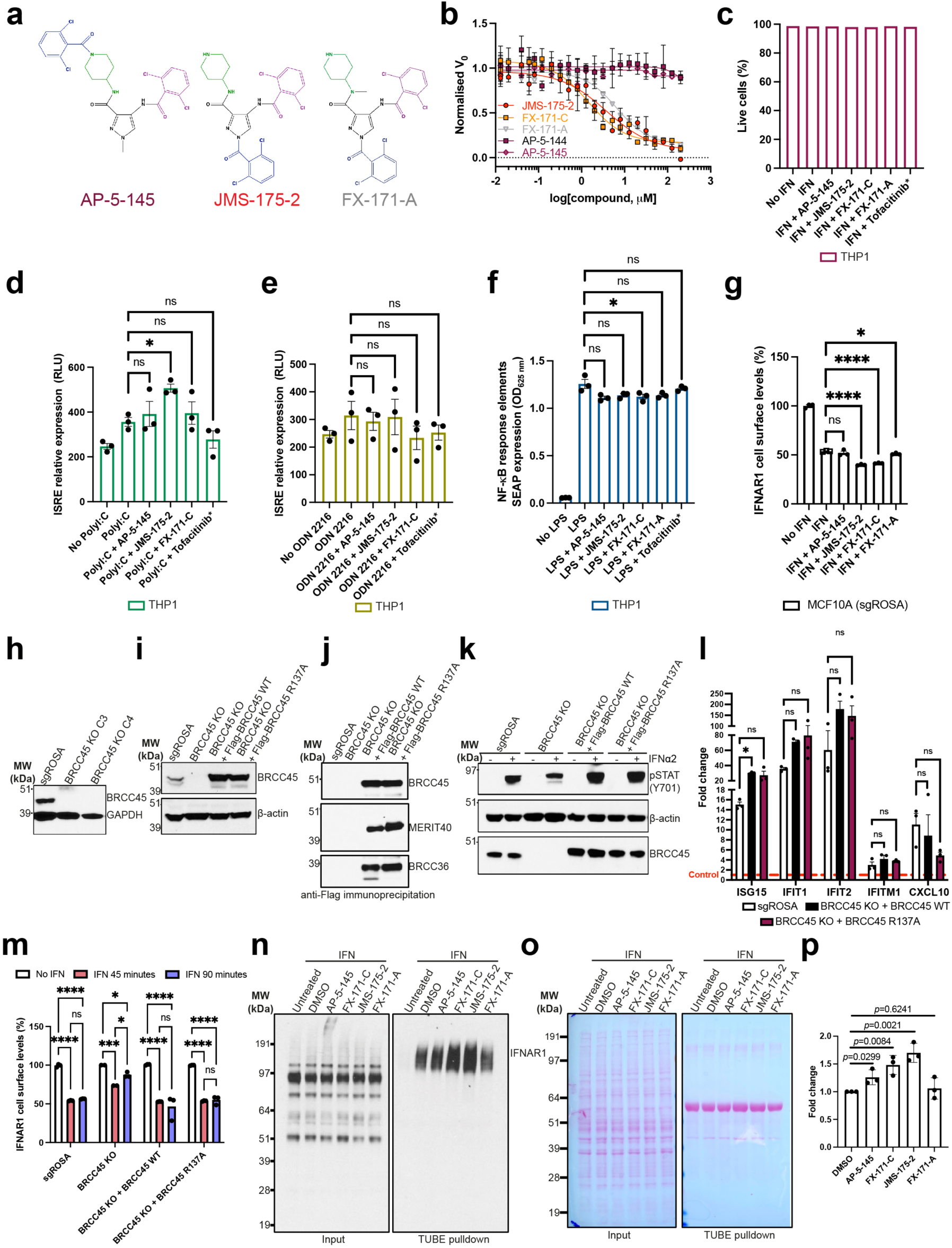
Establishing the effect of BLUE compound treatment on immune signalling pathways, IFNAR1 surface levels, and IFNAR1 ubiquitylation. **a,** Chemical structures of compounds AP-5-145, JMS-175-2, and FX-171-A. **b,** Dose-response inhibition of BRISC by indicated compounds. Data points are mean ± SEM for three independent experiments. **c,** Bar chart representing percentage of live cells across all conditions for ISRE expression and FACS analysis in THP-1 cells shown in **Figs. 4a, h**. THP-1 cells were treated with/without hIFNα2 (25 ng/mL) and either 4 μM inhibitor (JMS-175-2, FX-171-C, FX-171-A), 4 μM AP-5-145 negative control, DMSO control (0.1%), or JAK/STAT inhibitor Tofacitinib (*0.4 μM) for 16 hours. Bars represent the means from three independent experiments. **d, e,** Luciferase analysis of the ISRE in THP-1 supernatant after stimulation with **d,** polyI:C (1 µg/mL), or **e,** ODN 2216 (1 µM) and treatment with either 4 μM inhibitor (JMS-175-2, FX-171-C, FX-171-A), 4 μM AP-5-145 negative control, or JAK/STAT inhibitor Tofacitinib (*0.4 μM) for 16 hours. **f,** NF-κB pathway activity analysed by SEAP activity in THP-1 supernatants. Optical density measured at 625 nm. Data points in **d-f** are from three independent experiments. **g,** SgROSA MCF10A cells were treated with/without hIFN-Iα (50 ng/mL) and either 5 μM inhibitor (JMS-175-2, FX-171-C, FX-171-A), 5 μM negative control AP-5-145 or DMSO (0.1%) for 90 minutes. In **g, m,** IFNAR1 cell surface levels (%) were quantified using FACS analysis and calculated as a percentage of no IFN stimulation, and data points are from three independent experiments. **h,** BRCC45 protein levels in selected clones after knock out in MCF10A *Cas*9 cells. sgROSA was used as a CRISPR *Cas*9 control. **i,** BRCC45 expression in whole cell lysates from MCF10A *Cas*9 cells expressing CRISPR control (sgROSA), BRCC45 WT, and BRCC45 R137A. **j,** Anti-Flag co-immunoprecipitation performed in indicated MCF10A cell lines. BRISC complex subunits were detected using specific antibodies. **k,** MCF10A cell lines were treated with and without hIFNα2 (75 ng/mL) for 1 hour. STAT1 Tyr701 phosphorylation, BRCC45 and total protein levels (β-actin) were detected using specific antibodies. **l,** MCF10A *Cas*9 cells were treated with /without hIFNα2 for 4 hours. Expression of interferon-induced genes, ISG15, IFIT1, IFIT2, IFITM1, and CXCL10 were normalised to 18s rRNA and presented as fold change to own no IFN treated control. Data points are from three independent experiments. **m,** MCF10A cells (sgROSA, BRCC45 KO, BRCC45 WT and BRCC45 R137A) were treated with/without hIFN-Iα (50 ng/mL) for either 45 or 90 minutes. **n,** Anti-IFNAR1 immunoblots of TUBE-pulldown. *Left,* input samples after stimulation with IFNα2, treatment with BLUE inhibitors and cell lysis. *Right,* ubiquitylated IFNAR1 isolated with agarose-TUBE beads after IFNα2 stimulation. The Western blot shown is representative of three biological replicates. For raw, uncropped Western blots, see **Supplementary** Figure 1**. o,** Ponceau stained membranes of blots shown in **n**, prior to antibody incubation. **p,** Densitometry quantification of Western blot shown in **n,** and two other biological replicates (n=3). In **d-f,** paired t-tests were used to compare compound treated cells with DMSO control cells. In **g,** unpaired t-tests were used to compare compound treated cells with DMSO control cells. In **l,** a two-way ANOVA was used to gene expression levels for five genes in both BRCC45 WT and BRCC45 R137A cells to the sgROSA MCF10A cells. In **m,** a two-way ANOVA was performed to compare no IFN vs. IFN 45 minutes vs. IFN 90 minutes. P values illustrated by * <0.05, ** <0.01, *** <0.005, **** <0.0001, ns = non-significant. Error bars represent ± SEM.

**Extended Data Figure 9.**
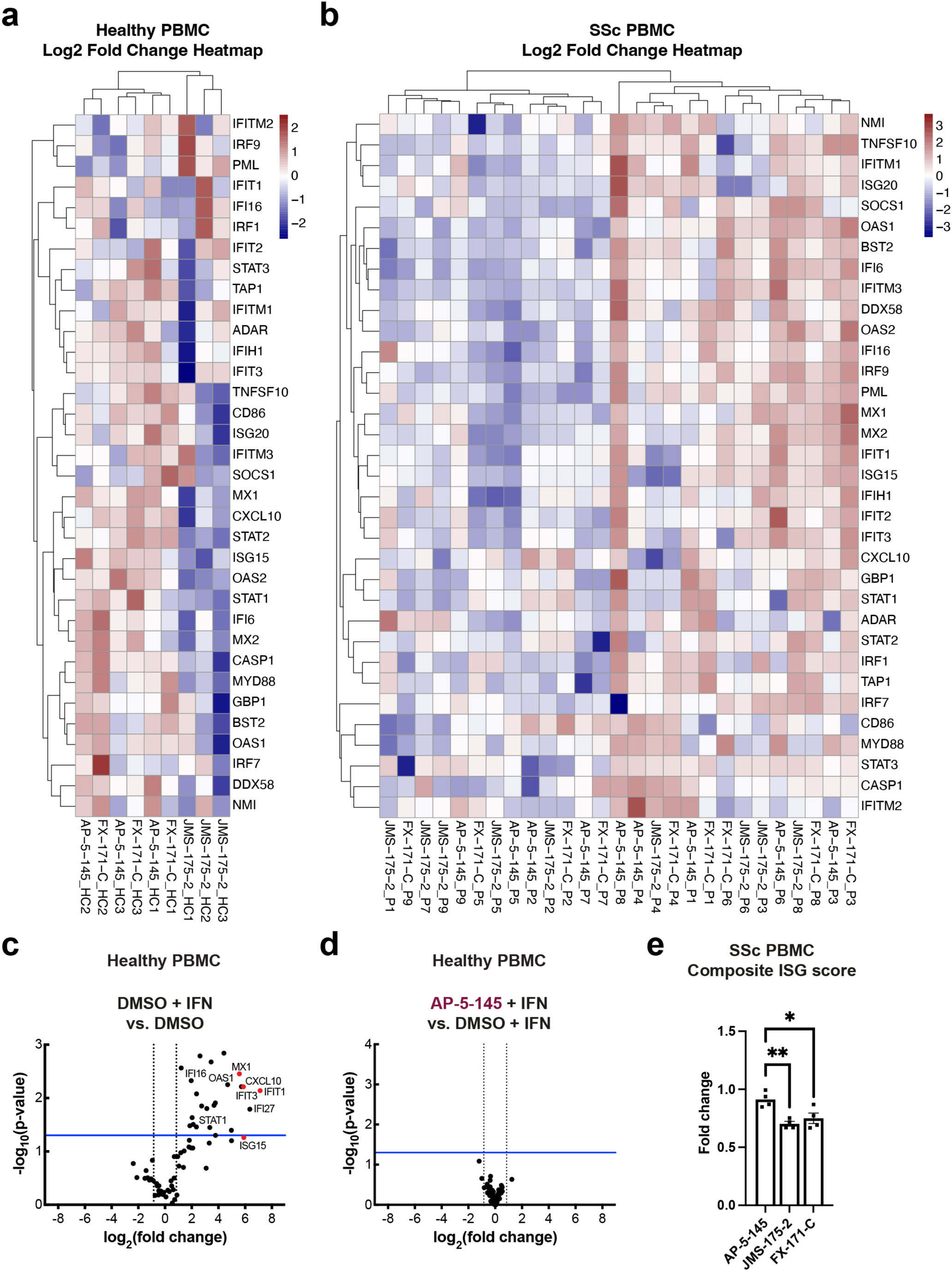
BLUE compounds reduce interferon-stimulated gene expression in stimulated healthy and unstimulated SSc PBMCs. **a, c, d,** Type I IFN signalling gene expression analysis of healthy control PBMCs treated with/without IFNα2 (20 ng/mL) and DMSO control (0.1 %) or 2 µM AP-5-145, JMS-175-2, or FX-171-C for 16 hours (n=3). **a,** Heatmap of each ISG expression levels relative to each donors housekeeping gene expression levels (geomean of ACTB, GAPDH, HPRT1, RPLP0), shown as Log2 fold change to grouped AP-5-145. Data shown for each individual donor, HC = healthy control. Heat map represents the mean fold change from three healthy donors.**c,** Volcano plot illustrating genes increased with addition of IFN + DMSO vs. DMSO only. **d,** Volcano plot illustrating no change in gene expression with negative control AP-5-145 + IFN vs. DMSO + IFN only. In **c,** and **d,** data points are the means from three independent experiments. **b,** As in **a,** Type I IFN signalling gene expression analysis of unstimulated SSc PBMCs from nine patients, treated with 2 µM AP-5-145, JMS-175-2, or FX-171-C for 16 hours. ISG relative expression to each donors housekeeping genes, shown as Log2 fold change relative to grouped AP-5-145, as in **a.** P refers to patient number i.e. P1 = patient 1. Heat map represents the mean fold change from nine SSc donors. **e,** PBMCs were isolated from patients and treated with DMSO (0.1%), 2 µM AP-5-145, FX-171-C or JMS-175-2 for 16 hours without IFN stimulation. Composite ISG score (including CXCL10, IFIT1, ISG15 and MX1) gene expression analysis between conditions relative to each donor DMSO control. Error bars represent ± SEM. Inividual data points represent the mean fold change for each gene for 20 donors.

**Extended Data Figure 10.**
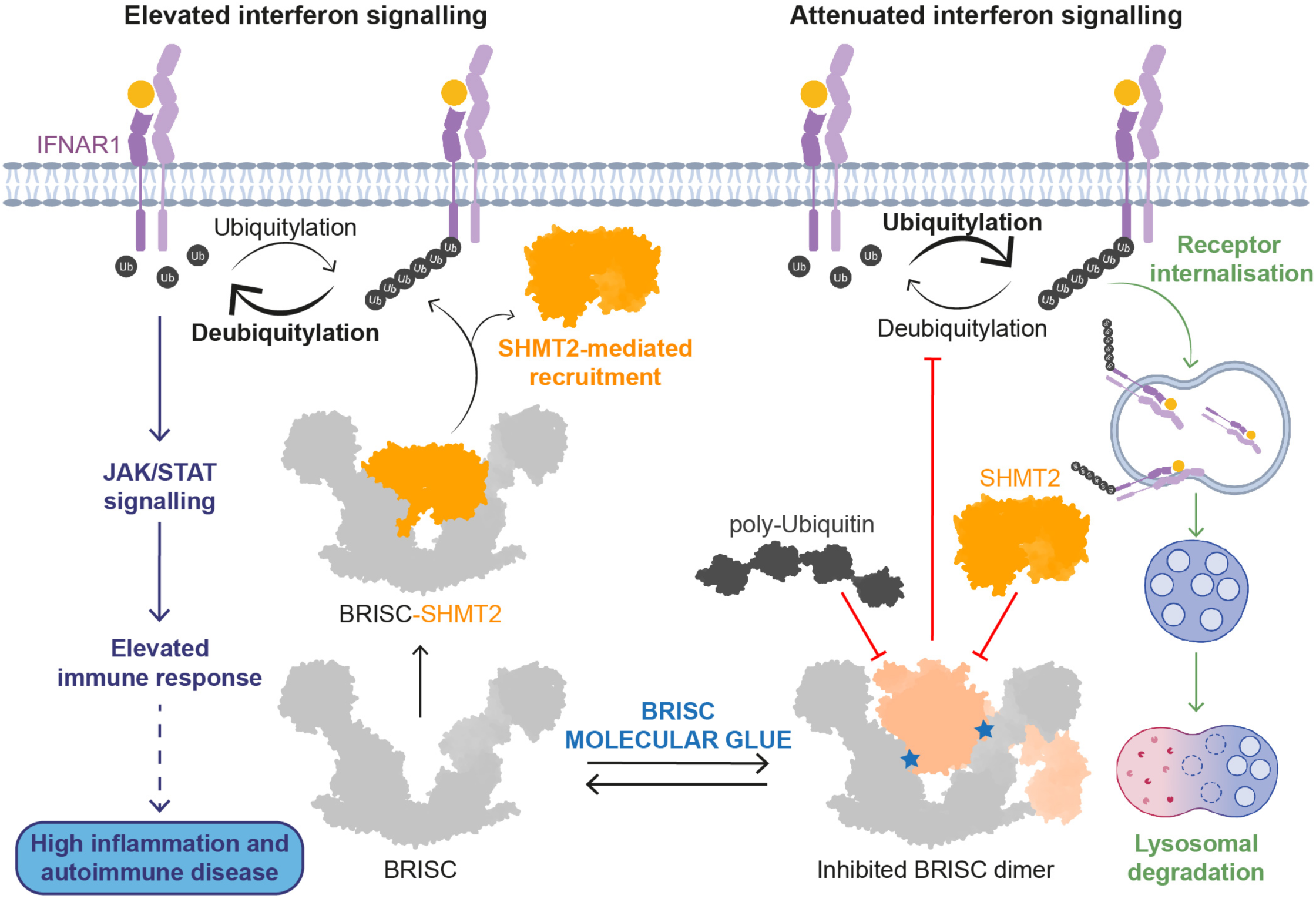
Proposed model of BLUE compound mode of action. Interferon binding to IFNAR1 receptors triggers JAK/STAT signalling and an elevated immune response. Interferon also initiates IFNAR1 receptor ubiquitylation (K63-linked), receptor internalisation and lysosomal degradation. The BRISC-SHMT2 complex is required for deubiquitylation of IFNAR1. BRISC is recruited to IFNAR1/2 through interactions with SHMT2 to promote sustained interferon signalling and inflammation. BLUE compounds (blue stars) promote formation of a BRISC dimer complex, which sterically hinders SHMT2 and polyubiquitin binding.

**Extended Data Table 1.** -Cryo-EM data collection, refinement, and validation statistics.

|  | **BRISCΔNΔC + FX-171-C**<br>EMD-17980, PDB 8PVY | **BRISCΔNΔC + JMS-175-2**<br>EMD-18009, PDB 8PY2 | **BRISC(FL)** |
| --- | --- | --- | --- |
| **Data collection and processing** |  |  |  |
| Microscope |  | FEI Titan Krios G2 |  |
| Detector | Thermo Scientific Falcon 4 | Thermo Scientific Falcon 4 | Thermo Scientific Falcon 4 |
| Energy filter | Thermo Scientific SelectrisX |  | Thermo Scientific SelectrisX |
| Energy filter slit (eV) | 10 |  |  |
| Magnification | 165,000 x | 96,000 x | 165,000 x |
| Voltage (kV) | 300 | 300 | 300 |
| Spot size | 7 | 6 | 8 |
| Illuminated area ($\mu\text{m}$) | 0.8 | 1 | 0.53 |
| Pixel size ($\text{\AA}$) | 0.71 | 0.82 | 0.74 |
| Defocus range ($\mu\text{m}$) | -1.6 to -2.5 | -1.7 to -3.1 | -0.9 to -2.7 |
| Total electron dose ($\text{e}^-/\text{\AA}^2$) | 34.97 | 39.84 | 40.46 |
| Exposure (s) | 3.43 | 5.99 | 2.78 |
| Number of frames | 44 | 40 | 40 |
| Electron dose per frame ($\text{e}^-/\text{\AA}^2$) | 0.8 | 0.99 | 1 |
| Movies collected | 16,750 | 7,768 | 14,573 |
| Acquisition mode | Counting | Counting | Counting |
| Symmetry imposed | C1 | C1 | C1, C2 |
| Initial particle images (no.) | 2,458,785 | 1,616,457 | 1,933,988 |
| Final particle images (no.) | 632,988 | 371,872 | 34,099 (monomer)<br>32,283 (dimer) |
| Map resolution ($\text{\AA}$) | 3.02 | 3.32 | 7.1 (monomer)<br>7.2 (dimer, C1)<br>8.3 (dimer, C2) |
| FSC threshold | 0.143 | 0.143 | 0.143 |
| Map resolution range ($\text{\AA}$) | 2.8 - 5.4 | 3.1 - 7.8 | 4.0 - 7.0 (monomer)<br>4.0 - 7.0 (dimer, C1)<br>5.0 - 9.0 (dimer, C1) |
| **Refinement** |  |  |  |
| Initial models used (PDB code) | 6R8F, 6H3C | 6R8F, 6H3C |  |
| Model resolution ($\text{\AA}$) | 2.7 | 3.1 | |
| FSC threshold | 0.143 | 0.143 |  |
| Model composition |  |  |  |
| Non-hydrogen atoms | 35736 | 35730 |  |
| Protein residues | 4436 | 4436 |  |
| Ligands | ZN: 4 | ZN: 4 |  |
|  | FX-171-C: 2 | JMS-175-2: 2 |  |
| *B* factors ($\text{\AA}^2$) | | | |
| Protein | 60.66 | 22.53 |  |
| Ligands | 18.46 | 11.95 |  |
| R.M.S.D. from ideal geometry |  |  |  |
| Bond lengths ($\text{\AA}$) | 0.003 | 0.002 | |
| Bond angles ($^\circ$) | 0.72 | 0.519 | |
| Validation |  |  |  |
| MolProbity score | 1.97 | 1.94 |  |
| Clashscore | 9.94 | 11.14 |  |
| Ramachandran plot |  |  |  |
| Favoured (%) | 92.85 | 94.47 |  |
| Allowed (%) | 7.15 | 5.53 |  |
| Disallowed (%) | 0 | 0 |  |
| Rotamer outliers (%) | 0.13 | 0.33 |  |
| Cβ outliers (%) | 0 | 0 | |
| Cis proline (%) | 0 | 0 |  |
| CaBLAM outliers (%) | 3.95 | 3.31 |  |

**Extended Data Table 2.** -HDX-MS Data Summary Table.

| **Data Set** | **BRISC + DMSO** | **BRISC + FX-171-C** |
| --- | --- | --- |
| **HDX reaction details** | 50 mM potassium phosphate, 200 mM NaCl, pD 7.5, 95 % D<sub>2</sub>O, 4°C | 50 mM potassium phosphate, 200 mM NaCl, pD 7.5, 95 % D<sub>2</sub>O, 4°C |
| **HDX time course (min)** | 0, 0.5, 1, 10, 60 |  |
| **HDX control samples** | Maximally-labelled controls were not performed. |  |
| **Back exchange** | Not applicable |  |
| **Number of peptides** | BRCC36: 110 | BRCC36: 111 |
|  | Abraxas2ΔC: 71 | Abraxas2ΔC: 71 |
|  | BRCC45: 110 | BRCC45: 109 |
|  | MERIT40ΔN: 89 | MERIT40ΔN: 89 |
| **Sequence coverage (%)** | BRCC36: 94.30 | BRCC36: 94.30 |
|  | Abraxas2ΔC: 86.89 | Abraxas2ΔC: 86.89 |
|  | BRCC45: 88.37 | BRCC45: 88.37 |
|  | MERIT40ΔN: 92.66 | MERIT40ΔN: 92.66 |
| **Average peptide length / Redundancy** | BRCC36: 9.87/3.64 | BRCC36: 9.88/3.68 |
|  | Abraxas2ΔC: 11.76/3.60 | Abraxas2ΔC: 11.76/3.60 |
|  | BRCC45: 9.99/3.21 | BRCC45: 10.00/3.19 |
|  | MERIT40ΔN: 9.21/3.42 | MERIT40ΔN: 9.21/3.42 |
| **Replicates (biological or technical)** | 3 technical | 3 technical |
| **Repeatability (average SD)** | BRCC36: 0.1185 | BRCC36: 0.0578 |
|  | Abraxas2ΔC: 0.1186 | Abraxas2ΔC: 0.0623 |
|  | BRCC45: 0.0861 | BRCC45: 0.0620 |
|  | MERIT40ΔN: 0.0927 | MERIT40ΔN: 0.0584 |
| **Significant differences in HDX (delta HDX > X D)** | Reference | 99% CI in summed data |

**Extended Data Table 3.** -Clinical details of patients included in the study.

| ID | LeRoy | WBC count<br>($\times 10^6$ cells/L) | Lymphocytes<br>($\times 10^6$ cells/L) | Low lymphocytes count | Neutrophils<br>($\times 10^6$ cells/L) | Low neutrophils count | Mtx | Mmf | Rtx | Overlap |
| --- | --- | --- | --- | --- | --- | --- | --- | --- | --- | --- |
| 4 | lcSSc | 5480 | 1700 | 0 | 2930 | 0 | 0 | 0 | 0 | 0 |
| 34 | lcSSc | 5210 | 1100 | 0 | 3670 | 0 | 0 | 0 | 1 | 0 |
| 36 | lcSSc | 7780 | 1440 | 0 | 5830 | 0 | 0 | 0 | 0 | 0 |
| 52 | lcSSc | 7280 | 2120 | 0 | 4160 | 0 | 0 | 0 | 0 | 0 |
| **68** | **lcSSc** | **5350** | **1510** | **0** | **3230** | **0** | **0** | **1** | **0** | **RA** |
| 74 | lcSSc | 10530 | 1440 | 0 | 8110 | 0 | 0 | 0 | 0 | 0 |
| **78** | **dcSSc** | **5000** | **1570** | **0** | **2700** | **0** | **0** | **0** | **0** | **0** |
| **79** | **lcSSc** | **8970** | **1350** | **0** | **6880** | **0** | **0** | **0** | **0** | **SjS** |
| 133 | lcSSc | 10240 | 1410 | 0 | 7920 | 0 | 0 | 0 | 0 | 0 |
| 134 | lcSSc | 4880 | 1349 | 0 | 2710 | 0 | 1 | 0 | 0 | RA |
| **142** | **lcSSc** | **8570** | **1530** | **0** | **6510** | **0** | **0** | **0** | **0** | **0** |
| 174 | lcSSc | 10160 | 850 | 1 | 8300 | 0 | 0 | 0 | 0 | SLE |
| 180 | lcSSc | 7710 | 2570 | 0 | 4220 | 0 | 0 | 0 | 0 | 0 |
| **205** | **lcSSc** | **8900** | **2880** | **0** | **4950** | **0** | **0** | **1** | **0** | **0** |
| **219** | **lcSSc** | **9800** | **1140** | **0** | **7300** | **0** | **0** | **0** | **1** | **0** |
| **244** | **lcSSc** | **7240** | **1600** | **0** | **4790** | **0** | **1** | **0** | **0** | **0** |
| 227 | lcSSc | 5960 | 1750 | 0 | 3710 | 0 | 0 | 0 | 0 | SjS |
| 266 | dcSSc | 8760 | 2130 | 0 | 5720 | 0 | 0 | 1 | 0 | 0 |
| 288 | lcSSc | 5710 | 1980 | 0 | 3210 | 0 | 0 | 1 | 0 | DM |
| 303 | lcSSc | 4710 | 1930 | 0 | 2960 | 0 | 0 | 0 | 0 | RA |
| 409 | lcSSc | 72800 | 2390 | 0 | 3990 | 0 | 1 | 0 | 0 | Chron's |

In superarray (n=9)
In CXCL10 ELISA (n=7)
LeRoy = LeRoy classification of Systemic Sclerosis, lcSSc = limited cutaneous Systemic Sclerosis, dcSSc = diffuse cutaneous Systemic Sclerosis. Normal range for WBC count: 4000-11000 $\times 10^6$ cells/L. Normal range for lymphocytes count: 1000-4500 $\times 10^6$ cells/L. Normal range for neutrophils count: 2000-7500 $\times 10^6$ cells/L. Mtx (Methotrexate), Mmf (Mycophenolate Mofetil), Rtx (Rituximab); 1 = yes, 0 = no. Overlap refers to secondary or overlap autoimmune conditions affecting the patients including: Rheumatoid Arthritis (RA), Systemic Lupus Erythematosus (SLE), Sjögren Syndrome (SjS), Chron's disease (Chron's), Dermatomyositis (DM).

## References

1. Hershko, A. & Ciechanover, A. The ubiquitin system. Annu. Rev. Biochem 67, 425–79 (1998).

2. Komander, D., Clague, M. J. & Urbé, S. Breaking the chains: Structure and function of the deubiquitinases. Nat Rev Mol Cell Biol 10, 550–563 (2009).

3. Komander, D. & Rape, M. The Ubiquitin Code. Annu Rev Biochem 81, 203–229 (2012).

4. Walden, M., Masandi, S. K., Pawłowski, K. & Zeqiraj, E. Pseudo-DUBs as allosteric activators and molecular scaffolds of protein complexes. Biochem Soc Trans 46, (2018).

5. Clague, M. J., Urbé, S. & Komander, D. Breaking the chains: deubiquitylating enzyme specificity begets function. Nat Rev Mol Cell Biol 20, 338–352 (2019).

6. Hu, H. & Sun, S. C. Ubiquitin signaling in immune responses. Cell Research vol. 26 457–483 Preprint at 10.1038/cr.2016.40 (2016).

7. Ross, C. A. & Pickart, C. M. The ubiquitin-proteasome pathway in Parkinson’s disease and other neurodegenerative diseases. Trends in Cell Biology vol. 14 703–711 Preprint at 10.1016/j.tcb.2004.10.006 (2004).

8. Sacco, J. J., Coulson, J. M., Clague, M. J. & Urbé, S. Emerging roles of deubiquitinases in cancer-associated pathways. IUBMB Life vol. 62 140–157 Preprint at 10.1002/iub.300 (2010).

9. Cohen, P. & Tcherpakov, M. Will the Ubiquitin System Furnish as Many Drug Targets as Protein Kinases? Cell vol. 143 686–693 (Cell Press, 2010).

10. Harrigan, J. A., Jacq, X., Martin, N. M. & Jackson, S. P. Deubiquitylating enzymes and drug discovery: Emerging opportunities. Nat Rev Drug Discov 17, 57–77 (2018).

11. Cooper, E. M. et al. K63-specific deubiquitination by two JAMM/MPN+ complexes: BRISC-associated Brcc36 and proteasomal Poh1. EMBO Journal 28, 621–631 (2009).

12. Feng, L., Wang, J. & Chen, J. The Lys63-specific deubiquitinating enzyme BRCC36 is regulated by two scaffold proteins localizing in different subcellular compartments. Journal of Biological Chemistry 285, 30982–30988 (2010).

13. Kim, H., Chen, J. & Yu, X. Ubiquitin-binding protein RAP80 mediates BRCA1-dependent DNA damage response. Science (1979) 316, 1202–1205 (2007).

14. Sobhian, B. et al. RAP80 targets BRCA1 to specific ubiquitin structures at DNA damage sites. Science (1979) 316, 1198–1202 (2007).

15. Wang, B. et al. Abraxas and RAP80 form a BRCA1 protein complex required for the DNA damage response. Science (1979) 316, 1194–1198 (2007).

16. Cooper, E. M., Boeke, J. D. & Cohen, R. E. Specificity of the BRISC deubiquitinating enzyme is not due to selective binding to Lys63-linked polyubiquitin. Journal of Biological Chemistry 285, 10344–10352 (2010).

17. Patterson-Fortin, J., Shao, G., Bretscher, H., Messick, T. E. & Greenberg, R. A. Differential regulation of JAMM domain deubiquitinating enzyme activity within the RAP80 complex. Journal of Biological Chemistry 285, 30971–30981 (2010).

18. Zeqiraj, E. et al. Higher-Order Assembly of BRCC36-KIAA0157 Is Required for DUB Activity and Biological Function. Mol Cell 59, 970–983 (2015).

19. Kyrieleis, O. J. P. et al. Three-Dimensional Architecture of the Human BRCA1-A Histone Deubiquitinase Core Complex. Cell Rep 17, 3099–3106 (2016).

20. Walden, M. et al. Metabolic control of BRISC–SHMT2 assembly regulates immune signalling. Nature 570, 194–199 (2019).

21. Rabl, J. et al. Structural Basis of BRCC36 Function in DNA Repair and Immune Regulation. Mol Cell 75, 483–497 (2019).

22. Jiang, Q. et al. Autologous K63 deubiquitylation within the BRCA1-A complex licenses DNA damage recognition. Journal of Cell Biology 221, (2022).

23. Zheng, H. et al. A BRISC-SHMT Complex Deubiquitinates IFNAR1 and Regulates Interferon Responses. Cell Rep 5, 180–193 (2013).

24. Pascual, V., Farkas, L. & Banchereau, J. Systemic lupus erythematosus: all roads lead to type I interferons. Current Opinion in Immunology vol. 18 676–682 Preprint at 10.1016/j.coi.2006.09.014 (2006).

25. Conigliaro, P. et al. The type I IFN system in rheumatoid arthritis. in Autoimmunity vol. 43 220–225 (Autoimmunity, 2010).

26. Wu, M. & Assassi, S. The role of type 1 interferon in systemic sclerosis. Frontiers in Immunology vol. 4 Preprint at 10.3389/fimmu.2013.00266 (2013).

27. Liang, Q. et al. A selective USP1-UAF1 inhibitor links deubiquitination to DNA damage responses. Nat Chem Biol 10, 298–304 (2014).

28. Turnbull, A. P. et al. Molecular basis of USP7 inhibition by selective small-molecule inhibitors. Nature 550, 481–486 (2017).

29. Kategaya, L. et al. USP7 small-molecule inhibitors interfere with ubiquitin binding. Nature 550, 534–538 (2017).

30. Yuan, T. et al. Inhibition of Ubiquitin-Specific Proteases as a Novel Anticancer Therapeutic Strategy. Frontiers in Pharmacology vol. 9 1080 Preprint at 10.3389/fphar.2018.01080 (2018).

31. Rennie, M. L., Arkinson, C., Chaugule, V. K. & Walden, H. Cryo-EM reveals a mechanism of USP1 inhibition through a cryptic binding site. Sci Adv 8, 6353 (2022).

32. Lauinger, L. et al. Thiolutin is a zinc chelator that inhibits the Rpn11 and other JAMM metalloproteases. Nat Chem Biol 13, 709–714 (2017).

33. Li, J. et al. Capzimin is a potent and specific inhibitor of proteasome isopeptidase Rpn11. Nat Chem Biol 13, 486–493 (2017).

34. Schlierf, A. et al. Targeted inhibition of the COP9 signalosome for treatment of cancer. Nat Commun 7, 1–10 (2016).

35. Lange, S. M., Armstrong, L. A. & Kulathu, Y. Deubiquitinases: From mechanisms to their inhibition by small molecules. Mol Cell 82, 15–29 (2022).

36. Schreiber, S. L. & Crabtree, G. R. The mechanism of action of cyclosporin A and FK506. Immunol Today 13, 136–142 (1992).

37. Tan, X. et al. Mechanism of auxin perception by the TIR1 ubiquitin ligase. Nature 446, 640–645 (2007).

38. Liu, J. et al. Calcineurin is a common target of cyclophilin-cyclosporin A and FKBP-FK506 complexes. Cell 66, 807–815 (1991).

39. Schreiber, S. L. The Rise of Molecular Glues. Cell 184, 3–9 (2021).

40. Ito, T. et al. Identification of a primary target of thalidomide teratogenicity. Science (1979) 327, 1345– 1350 (2010).

41. Wyatt, P. G. et al. Identification of N-(4-piperidinyl)-4-(2,6-dichlorobenzoylamino)-1H-pyrazole-3-carboxamide (AT7519), a novel cyclin dependent kinase inhibitor using fragment-based X-ray crystallography and structure based drug design. J Med Chem 51, 4986–4999 (2008).

42. Michel, M. A. et al. Assembly and Specific Recognition of K29-and K33-Linked Polyubiquitin. Mol Cell 58, 95–109 (2015).

43. McCullough, J., Clague, M. J. & Urbé, S. AMSH is an endosome-associated ubiquitin isopeptidase. Journal of Cell Biology 166, 487–492 (2004).

44. Sato, Y. et al. Structural basis for specific cleavage of Lys 63-linked polyubiquitin chains. Nature 455, 358– 362 (2008).

45. Li, J. et al. Epidithiodiketopiperazines Inhibit Protein Degradation by Targeting Proteasome Deubiquitinase Rpn11. Cell Chem Biol 25, 1350–1358.e9 (2018).

46. Hodge, C. D. et al. Covalent Inhibition of Ubc13 Affects Ubiquitin Signaling and Reveals Active Site Elements Important for Targeting HHS Public Access Author manuscript. ACS Chem Biol 10, 1718–1728 (2015).

47. Ceccarelli, D. F. et al. An allosteric inhibitor of the human Cdc34 ubiquitin-conjugating enzyme. Cell 145, 1075–1087 (2011).

48. Ross, R. L. et al. Targeting human plasmacytoid dendritic cells through BDCA2 prevents skin inflammation and fibrosis in a novel xenotransplant mouse model of scleroderma. Ann Rheum Dis 80, 920–929 (2021).

49. Fraile, J. M., Quesada, V., Rodríguez, D., Freije, J. M. P. & López-Otín, C. Deubiquitinases in cancer: New functions and therapeutic options. Oncogene 31, 2373–2388 (2012).

50. Liu, B. et al. Deubiquitinating enzymes (DUBs): decipher underlying basis of neurodegenerative diseases. Mol Psychiatry 27, 259–268 (2022).

51. Zinngrebe, J., Montinaro, A., Peltzer, N. & Walczak, H. Ubiquitin in the immune system. EMBO Rep 15, 28–45 (2014).

52. Kimani, S. W., et al. The co-crystal structure of Cbl-b and a small-molecule inhibitor reveals the mechanism of Cbl-b inhibition. Commun Biol 6, (2023).

53. Collins, G. P. et al. A First-in-Human Phase 1 Trial of NX-1607, a First-in-Class Oral CBL-B Inhibitor, in Patients with Advanced Malignancies Including DLBCL. Blood 142, 3093–3093 (2023).

54. Gersch, M. et al. Distinct USP25 and USP28 Oligomerization States Regulate Deubiquitinating Activity. Mol Cell 74, 436–451.e7 (2019).

55. Sauer, F. et al. Differential Oligomerization of the Deubiquitinases USP25 and USP28 Regulates Their Activities. Mol Cell 74, 421–435.e10 (2019).

56. Burkhart, R. A. et al. Mitoxantrone targets human ubiquitin-specific peptidase 11 (USP11) and is a potent inhibitor of pancreatic cancer cell survival. Molecular Cancer Research 11, 901–911 (2013).

57. Ward, S. J. et al. The structure of the deubiquitinase USP15 reveals a misaligned catalytic triad and an open ubiquitin-binding channel. Journal of Biological Chemistry 293, 17362–17374 (2018).

58. Kategaya, L. et al. USP7 small-molecule inhibitors interfere with ubiquitin binding. Nature 550, 534–538 (2017).

59. Griffith, J. P. et al. X-ray structure of calcineurin inhibited by the immunophilin-immunosuppressant FKBP12-FK506 complex. Cell 82, 507–522 (1995).

60. Huai, Q. et al. Crystal structure of calcineurin-cyclophilin-cyclosporin shows common but distinct recognition of immunophilin-drug complexes. Proc Natl Acad Sci U S A 99, 12037–12042 (2002).

61. Cao, S. et al. Defining molecular glues with a dual-nanobody cannabidiol sensor. Nature Communications 2022 13:1 13, 1–14 (2022).

62. Fitzgerald, D. J. et al. Protein complex expression by using multigene baculoviral vectors. Nat Methods 3, 1021–1032 (2006).

63. Sonn-Segev, A. et al. Quantifying the heterogeneity of macromolecular machines by mass photometry. Nat Commun 11, (2020).

64. Zivanov, J. et al. New tools for automated high-resolution cryo-EM structure determination in RELION-3. Elife 7, (2018).

65. Zivanov, J., Nakane, T. & Scheres, S. H. W. Estimation of high-order aberrations and anisotropic magnification from cryo-EM data sets in RELION-3.1. IUCrJ 7, 253–267 (2020).

66. Nakane, T. et al. Single-particle cryo-EM at atomic resolution. Nature 587, 152–156 (2020).

67. Rohou, A. & Grigorieff, N. CTFFIND4: Fast and accurate defocus estimation from electron micrographs. J Struct Biol 192, 216–221 (2015).

68. Zhang, K. Gctf: Real-time CTF determination and correction. J Struct Biol 193, 1–12 (2016).

69. Scarff, C. A., Fuller, M. J. G., Thompson, R. F. & Iadaza, M. G. Variations on negative stain electron microscopy methods: Tools for tackling challenging systems. Journal of Visualized Experiments 2018, (2018).

70. Wagner, T. et al. SPHIRE-crYOLO is a fast and accurate fully automated particle picker for cryo-EM. Communications Biology 2019 2:1 2, 1–13 (2019).

71. Punjani, A., Rubinstein, J. L., Fleet, D. J. & Brubaker, M. A. CryoSPARC: Algorithms for rapid unsupervised cryo-EM structure determination. Nat Methods 14, 290–296 (2017).

72. Punjani, A., Zhang, H. & Fleet, D. J. Non-uniform refinement: adaptive regularization improves single-particle cryo-EM reconstruction. Nat Methods 17, 1214–1221 (2020).

73. Pettersen, E. F. et al. UCSF ChimeraX: Structure visualization for researchers, educators, and developers. Protein Science 30, 70–82 (2021).

74. Asarnow, D., Palovcak, E. & Cheng, Y. UCSF pyem v0.5. Zenodo 10.5281/zenodo.3576630. Preprint at https://doi.org/10.5281/zenodo.3576630 (2019).

75. Pettersen, E. F. et al. UCSF Chimera--a visualization system for exploratory research and analysis. J Comput Chem 25, 1605–1612 (2004).

76. Emsley, P., Lohkamp, B., Scott, W. G. & Cowtan, K. Features and development of Coot. Acta Crystallogr D Biol Crystallogr 66, 486–501 (2010).

77. Casañal, A., Lohkamp, B. & Emsley, P. Current developments in Coot for macromolecular model building of Electron Cryo-microscopy and Crystallographic Data. Protein Science 29, 1069–1078 (2020).

78. Schüttelkopf, A. W. & Van Aalten, D. M. F. PRODRG: A tool for high-throughput crystallography of protein-ligand complexes. Acta Crystallogr D Biol Crystallogr 60, 1355–1363 (2004).

79. Moriarty, N. W., Grosse-Kunstleve, R. W. & Adams, P. D. Electronic ligand builder and optimization workbench (eLBOW): A tool for ligand coordinate and restraint generation. Acta Crystallogr D Biol Crystallogr 65, 1074–1080 (2009).

80. Adams, P. D. et al. PHENIX: A comprehensive Python-based system for macromolecular structure solution. Acta Crystallogr D Biol Crystallogr 66, 213–221 (2010).

81. Edgar, R. C. MUSCLE: Multiple sequence alignment with high accuracy and high throughput. Nucleic Acids Res 32, 1792–1797 (2004).

82. Bond, C. S. & Schüttelkopf, A. W. ALINE: A WYSIWYG protein-sequence alignment editor for publication-quality alignments. Acta Crystallogr D Biol Crystallogr 65, 510–512 (2009).

83. Marty, M. T. et al. Bayesian deconvolution of mass and ion mobility spectra: From binary interactions to polydisperse ensembles. Anal Chem 87, 4370–4376 (2015).

84. Lau, A. M., Claesen, J., Hansen, K. & Politis, A. Deuteros 2.0: Peptide-level significance testing of data from hydrogen deuterium exchange mass spectrometry. Bioinformatics 37, 270–272 (2021).

85. Masson, G. R. et al. Recommendations for performing, interpreting and reporting hydrogen deuterium exchange mass spectrometry (HDX-MS) experiments. Nat Methods 16, 595–602 (2019).

